# Targeted and random mutagenesis of cassava brown streak disease susceptibility factors reveal molecular determinants of disease severity

**DOI:** 10.1101/2025.02.27.640518

**Authors:** ZJ Daniel Lin, Myia K. Stanton, Gabriela L. Hernandez, Elizabeth J. De Meyer, Zachary von Behren, Katherine Benza, Helene Tiley, Anna Meirink, Emerald Hood, Greg Jensen, Kerrigan B. Gilbert, James C. Carrington, Rebecca S. Bart

## Abstract

Cassava brown streak disease (CBSD) is caused by cassava brown streak viruses (CBSVs) from the family *Potyviridae*. Potyvirid viral genome-linked protein (VPg) recruitment of host eukaryotic translation initiation factor 4E (eIF4E) proteins is a critical step in the viral life cycle. CBSV VPg interacts with all five cassava eIF4E-family members. Simultaneously knocking out *eIF4E-*family genes *nCBP-1* and *nCBP-2*, in cultivar 60444, strongly reduces CBSD root symptoms and viral titer but does not result in complete resistance, likely due to gene family redundancy. To test for redundancy, we generated single and double mutants for each clade of the *eIF4E* gene family in farmer preferred cultivar TME419. Double mutants for the *eIF(iso)4E* and *nCBP* clades both exhibited reduced symptom severity, with *ncbp-1 ncbp-2* having the strongest phenotype. A yeast two-hybrid screen for nCBP-2 mutants that lose VPg affinity identified fifty-one mutants, including an L51F mutant. This finding is consistent with one of the recovered cassava mutants that had a 6 amino acid deletion, including L51, in nCBP-2 and showed a reduction in symptoms relative to wild type. The data presented here suggest that generating mutations corresponding to L51F of nCBP-2 in multiple or all five cassava eIF4E proteins may lead to stronger resistance to CBSD while avoiding pleiotropic effects.

## Introduction

Cassava is one of the most important starch crops grown in the world (*Agricultural production statistics 2000–2022*, 2023). In sub-Saharan Africa, cassava is particularly important for food security, but its production is threatened by viral diseases (Rey & Vanderschuren, 2017). In particular, cassava brown streak virus (CBSV) and Ugandan cassava brown streak virus (UCBSV), closely related ipomoviruses belonging to the family *Potyviridae*, are of major concern (Tomlinson *et al*., 2018; Robson *et al*., 2023). Independent or simultaneous infection by these viruses cause cassava brown streak disease (CBSD). Symptoms of CBSD include feathery chlorosis of leaves, brown streaking of stems, and necrosis of storage roots. Cassava varieties with varying levels of resistance have been identified and are being deployed, but none of the varieties currently available to farmers are fully immune (Sheat & Winter, 2023; Robson *et al*., 2023). A highly effective RNAi-based transgenic approach has been developed but faces regulatory hurdles in many target countries (Odipio *et al*., 2014; Beyene *et al*., 2016; Wagaba *et al*., 2016; Tomlinson *et al*., 2018).

Viruses of *Potyviridae* are positive sense, single stranded RNA viruses (Valli *et al*., 2015). Many of these viruses have been studied extensively as a large number are agronomically important pests (Revers & García, 2015; Yang *et al*., 2021). In many potyvirus-plant pathosystems, members of the host *eIF4E* gene family function as susceptibility factors and are necessary for completion of the viral life cycle (Robaglia & Caranta, 2006; Naderpour *et al*., 2010; Hart & Griffiths, 2013; Bastet *et al*., 2017; Gomez *et al*., 2019; Gao *et al*., 2020). This gene family encodes mRNA cap-binding proteins that facilitate recruitment of translation machinery to transcripts (Browning & Bailey-Serres, 2015). The plant *eIF4E-*family can be broken into three clades: *eIF4E*, *eIF(iso)4E*, and *nCBP* (Gomez *et al*., 2019). Previous work in *Arabidopsis* demonstrated that turnip mosaic virus (TuMV) requires a functional copy of *eIF(iso)4E* for disease (Duprat *et al*., 2002). Since then, potyvirus resistance loci in many crops have been mapped to *eIF4E-*family genes and alleles conferring potyvirus resistance often harbor missense mutations (Bastet *et al*., 2017). Additional studies have demonstrated that potyvirid viral protein genome-linked (VPg) interacts with eIF4E-family proteins (Wittmann *et al*., 1997; Schaad *et al*., 2000) confirming that the interacting *eIF4E-*family member functions as a susceptibility factor (Yeam *et al*., 2007; Charron *et al*., 2008). Conversely, viral isolates that break *eIF4E*-family based resistance have been found to have accumulated compensating mutations in their VPg (Charron *et al*., 2008; Gallois *et al*., 2010; Li *et al*., 2016; Lebaron *et al*., 2016; Takakura *et al*., 2018). VPg is covalently linked to the 5’ end of the viral genome. It has been hypothesized that VPg interacts with eIF4E-family proteins to recruit translational machinery to synthesize the viral polypeptide (Sanfaçon, 2015) and complete the viral life cycle. Solving the structure of VPg-susceptibility factor protein complex could facilitate engineering of susceptibility factor alleles that cannot be co-opted by the pathogen. However, co-crystal structures have yet to be produced and the VPg protein complex with eIF4E-family *S* factors falls below the current minimum size limit required for cryogenic electron microscopy. To date, engineering synthetic *eIF4E* resistance alleles has involved transferring resistance-conferring mutations from one pathosystem to another with some degree of success (Kim *et al*., 2014; Bastet *et al*., 2018, 2019; Zafirov *et al*., 2021).

We previously found that simultaneous mutations in two cassava eIF4E-family members, *nCBP-1* and *nCBP-2,* strongly attenuated CBSD storage root necrosis and accumulation of CBSV in storage roots of cultivar 60444 (Gomez *et al*., 2019). Knocking out *nCBP-2* alone resulted in an intermediate phenotype, while the *ncbp-1* single mutant exhibited no differences from wild type. Aerial symptoms persisted in the *ncbp-1 ncbp-2* double mutant, although with less severe stem brown streaking. This incomplete resistance suggests that VPg may interact broadly with the remaining cassava eIF4E-family proteins. Indeed, CBSV VPg could associate with all five cassava eIF4E-family proteins by coimmunoprecipitation (Gomez *et al*., 2019). As full CBSD resistance may require engineering all cassava *eIF4E*-family alleles to evade CBSV VPg, we employed targeted and random mutagenesis of nCBP-2 to identify structural determinants of VPg interaction. Furthermore, to better understand the degree of functional redundancy exhibited by cassava eIF4E-family proteins in CBSD, we here expand our CRISPR/Cas enabled genetic approach to characterize the roles of cassava eIF4E and eIF(iso)4E clades in CBSD.

## Results

### Generation of cassava *eIF4E-*family mutants

We previously reported that a cassava *ncbp-1 ncbp-2* double mutant in the 60444 background displayed increased, but incomplete, resistance to CBSV challenge (Taylor *et al*., 2012; Gomez *et al*., 2019). While VPg interacted most strongly with nCBP-1 and nCBP-2 in yeast two-hybrid assays, all five eIF4E-family proteins interacted with VPg in co-immunoprecipitation assays. These results suggest that there is likely functional redundancy among this family of proteins. To test this idea, we used CRISPR/Cas9-mediated genome editing to generate new cassava *eIF4E*-family mutants in TME419, a farmer preferred cassava cultivar. gRNAs were designed to target the first exon of each target gene and cassava transformations were performed as previously described (Fig. S2) (Taylor *et al*., 2012; Gomez *et al*., 2019). A total of 156 recovered lines from 21 transformations were characterized for possible mutations at the target site(s) using amplicon sequencing. These efforts yielded independent single mutant lines for all members of the cassava *eIF4E* gene family, along with *eif(iso)4e-1 eif(iso)4e-2* and *ncbp-1 ncbp-2* double mutants (Table 1, S3). All but one mutant line used in this study are predicted to have frameshifting mutations that result in loss of protein function. The lone exception is one of the *ncbp-1 ncbp-2* double mutants where both *nCBP-1* alleles and one allele of *nCBP-2* have frameshifting mutations and the remaining *nCBP-2* allele has a deletion that results in loss of amino acids 45 through 51. This allele is designated *nCBP-2^K45_L51del^*and the mutant line is designated *ncbp-1 nCBP-2^K45_L51del^* (Table 1, S3). No differences in growth and development were observed between any of the mutant lines and wild type (Fig. S3-4).

**Table 1.** TME419 CRISPR mutant genotypes.

| Line # | CRISPR target | Gene/allele | INDEL | Protein nomenclature |
| --- | --- | --- | --- | --- |
| 6-95 | *eIF4E* | *A* | -2 | S21*fsX1 |
|  |  | *B* | – | " |
| 172 | *iso4E1+2* | *iso4E1 A* | -1 | A11PfsX17 |
|  |  | *iso4E1 B* | -2 | A11HfsX9 |
|  |  | *iso4E2 A* | -2 | I7RfsX29 |
|  |  | *iso4E2 B* | 1 | E8RfsX36 |
| 92 | *iso4E1+2* | *iso4E1 A* | -2 | A11HfsX9 |
|  |  | *iso4E1 B* | -1 | A11PfsX17 |
|  |  | *iso4E2 A* | -2 | I7RfsX29 |
|  |  | *iso4E2 B* | 1 | E8RfsX36 |
| 10-9 | *iso4E2* | *A* | 1 | E8RfsX36 |
|  |  | *B* | – | " |
| 3-29 | *iso4E1* | *A* | -1 | A11PfsX17 |
|  |  | *B* | – | " |
| 93 | *nCBP1* | *A* | 1 | S23YfsX4 |
|  |  | *B* | -2 | S23TfsX3 |
| 4-53 | *nCBP2* | *A* | 1 | T23NfsX8 |
|  |  | *B* | – | " |
| 213 | *nCBP1+2* | *nCBP1 A* | 1 | S23YfsX4 |
|  |  | *nCBP1 B* | – | " |
|  |  | *nCBP2 A* | -4 | T22HfsX10 |
|  |  | *nCBP2 B* | -2 | T23IfsX7 |
| 40 | *nCBP1+2* | *nCBP1 A* | 1 | S23YfsX4 |
|  |  | *nCBP1 B* | 1 | P24TfsX3 |
|  |  | *nCBP2 A* | -21 | K45_L51del |
|  |  | *nCBP2 B* | -1 | A46LfsX31 |

### CBSD phenotypes of TME419 *eIF4E*-family mutants

To determine the contribution of cassava *eIF4E* family genes to development of CBSD, mutant and wild type TME419 were challenged with CBSV isolate Naliendele in greenhouse trials where stem brown streaking and leaf feathery chlorosis were tracked for approximately three months. The specific CBSV isolate was chosen as it produces strong aerial symptoms quickly after inoculation and reliably results in strong storage root necrosis (Wagaba *et al*., 2013). At the end of the viral challenges, storage roots were excavated from soil, sectioned, and scored for severity of necrosis as a measure of spread along storage root length and diameter (Gomez *et al*., 2019). These same root samples were used to measure viral titer by quantitative real-time PCR (qRT-PCR). For aboveground symptoms, *eif4e*, *eif(iso)4e-1, eif(iso)4e-2,* and *ncbp-1* single mutants had no reduction in disease severity relative to wild type, while *ncbp-*2 and both *eif(iso)4e* and *ncbp* double mutants exhibited small but significant reductions (Figure 1a). For storage root necrosis symptoms in the single mutants, the *ncbp-1* and *ncbp-2* mutants displayed a decrease in symptoms compared to wild type with the latter showing the strongest phenotype (Figure 1b). Each of the double mutants showed a decrease in symptom with the *ncbp-1 ncbp-2* mutant, where all *nCBP* alleles have frameshifting mutations, having the greatest reduction in necrosis; virtually no necrosis was observed (Figure 1b, d, S5_). We further measured viral accumulation in storage roots of wild type, *ncbp-1*, *ncbp-2*, *ncbp-1 ncbp-2*, and *ncbp-1 nCBP-2^K45_L51del^* lines using qPCR analysis and observed that lines with frameshifting mutations in *ncbp-2* had significantly lower virus titer (Figure 1c). Taken together, these data suggest that while nCBP-2 is the primary contributor to root disease severity, other eIF4E-family members may also contribute throughout the plant.

**Fig. 1.**
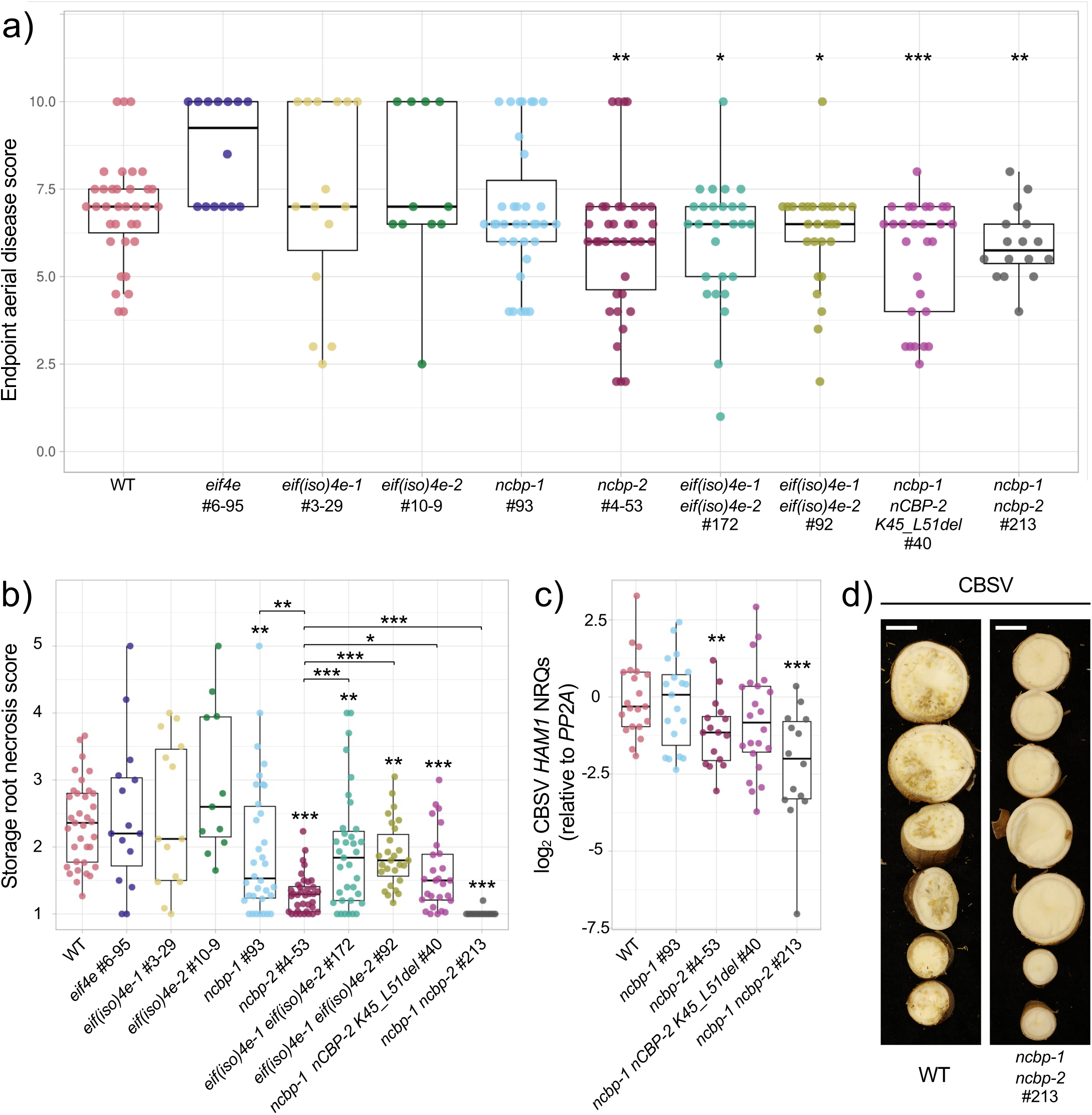
Endpoint CBSD storage root disease severity in cassava eIF4E-family mutants. a, b) Endpoint aerial disease (a) and storage root necrosis (b) observed in cassava eIF4E-family single and double mutants after challenge with CBSV isolate Naliendele. Data presented are an aggregate from all experimental replicates. Leaf and stem disease symptoms are evaluated separately on a 0-5 scale and then added together for a total aerial symptom score. Storage root necrosis severity was scored on a 1 to 5 scale, 1 being asymptomatic and 5 corresponding to severe necrosis throughout all storage roots of a plant. Mann-Whitney U test was used to detect statistical differences with an alpha = 0.05. * ≤ 0.05, ** ≤ 0.01, *** ≤ 0.001. Asterisks above boxplots denote differences from wild type, while asterisks above brackets indicate differences between indicated genotypes. c) qRT-PCR quantification of storage root CBSV viral levels from challenged mutants. CBSV HAM1 was quantified relative to cassava PP2A to obtain HAM1 normalized relative quantities (NRQs). HAM1 NRQs for each genotype are presented as relative to the mean of wild type. Statistical differences were detected with the Mann-Whitney U test as described in (a, b). d) Representative storage root sections from wild type and ncbp-1 ncbp-2 #213 plants challenged with CBSV. Scale bar denotes 1 cm.

### Purified CBSV VPg and cassava eIF4E-family proteins interact *in vitro*

To further explore functional redundancy within the eIF4E-family with respect to CBSD and, specifically, interaction with VPg, we used a biochemical approach. Using co-immunoprecipitation, we previously demonstrated that CBSV VPg and cassava eIF4E-family proteins form complexes in whole cell lysates using co-immunoprecipitation (Gomez *et al*., 2019). However, these experiments did not address whether the proteins are directly interacting or if another complex member is present and facilitating the protein-protein association. Thus, we tested if VPg directly associates with eIF4E-family proteins in the absence of additional factors using a GST pull-down experiment. GST-tagged cassava eIF4E-family proteins and VPg, tagged with both 6xHis and 3xFLAG, were expressed in and purified from *E. coli* (Figure S6). We found that CBSV VPg was pulled down by GST-tagged eIF4E, nCBP-1, and nCBP-2 in comparison to GST alone (Figure 2a, b). Pulldown of VPg by GST-tagged eIF(iso)4E-1 and eIF(iso)4E-2 was also tested but we observed inconsistent enrichment of VPg by GST-eIF(iso)4E-1 & −2, relative to GST alone, across experimental replicates.

**Figure 2.**
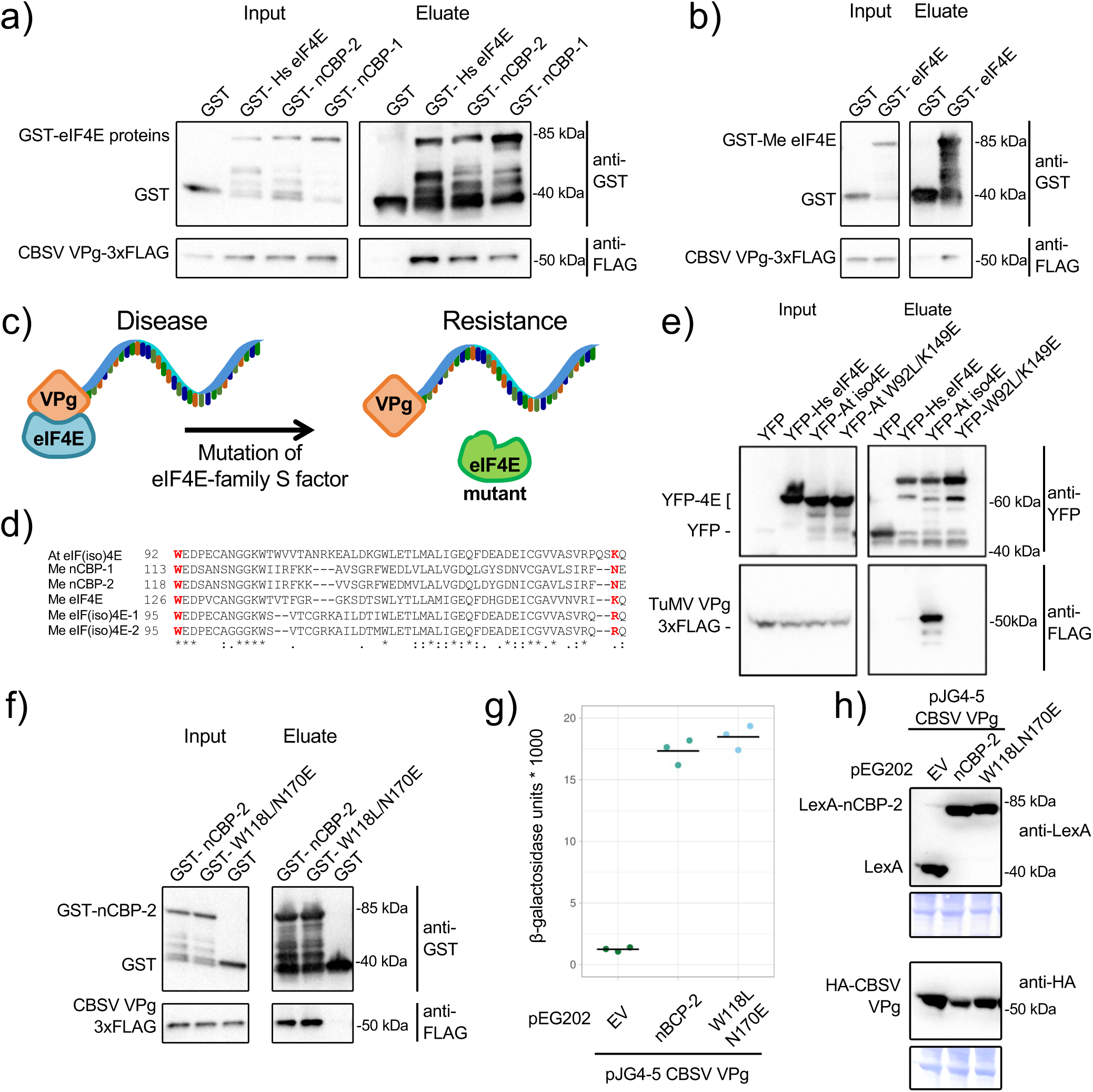
Interaction of GST-cassava eIF4E family proteins and CBSV VPg *in vitro*. a, b) GST-pulldown assays were performed with CBSV VPg-6xHis-3xFLAG and GST or GST-tagged cassava eIF4E-family proteins. Proteins were expressed in and purified from *E. coli*. Human (Hs) eIF4E was also tested in (a). c) Model for eIF4E-mediated resistance against potyvirids. Cartoon depicts potyvirid genome-linked VPg protein bound to eIF4E-family protein in order to cause disease, while mutations that disrupt this interaction results in resistance. d) Coimmunoprecipitation of YFP or YFP-eIF4E-family proteins with TuMV 6xHis-VPg-6xHIS-3xFLAG. VPg was expressed in and purified from *E. coli* and then mixed with extract from *N. benthamiana* leaves expressing YFP or YFP-fusion proteins. e) *Arabidopsis* eIF(iso)4E residues 92 through 150 aligned with homologous regions of cassava eIF4E-family proteins. W92 and K150, marked red, are necessary for interaction with TuMV VPg. * indicates conservation across all aligned proteins, : indicates conservation of amino acids with strongly similar properties, and . indicates conservation of amino acids with weakly similar properties. f) GST-pulldown assay as in a) and b) with CBSV VPg incubated with either GST, GST-nCBP-2, and GST-nCBP-2^W118L/N170E^. g) Comparison of nCBP-2 and nCBP-2^W118L/N170E^ interaction with CBSV VPg by quantitative yeast two-hybrid assay. h) Western blot analysis of nCBP-2 and VPg expression in yeast from g).

### Targeted nCBP-2 mutations fail to disrupt interaction with VPg

If CBSV VPg can coopt all cassava eIF4E-family proteins for completing the viral lifecycle, engineering cassava to be resistant to CBSV may require identifying alleles of *eIF4E*-family genes encoding mutations that disrupt VPg binding but still allow for translation-initiation activity. In other plant-potyvirid pathosystems, specific mutations within eIF4E-family members have proved sufficient to block interaction with VPg while maintaining normal protein function (Figure 2c) (Ashby *et al*., 2011; Kim *et al*., 2014). One well studied example is TuMV VPg and *Arabidopsis* eIF(iso)4E. In this system, mutation of a highly conserved tryptophan (W92L) and semi-conserved lysine (K149E) in *Arabidopsis* eIF(iso)4E disrupts binding of TuMV VPg in pull-down and co-immunoprecipitation experiments. Overexpression of this mutant *eIF(iso)4E* allele in Chinese cabbage confers resistance to TuMV (Miyoshi *et al*., 2006; Kim *et al*., 2014). To test the hypothesis that specific *eIF4E* gene family alleles might avoid VPg binding, we took a yeast two-hybrid approach using different alleles of *nCBP-2*. We recapitulated the W92L/K149E mutations from Arabidopsis eIF(iso)4E in cassava nCBP-2, resulting in nCBP-2^W118L/N170E^ (Figure 2d). Whereas the eIF(iso)4E^W92L/K149E^ no longer associated with TuMV VPg in a co-immunoprecipitation experiment (Figure 2e), we observed that CBSV VPg was similarly present in both wild type nCBP-2 and nCBP-2^W118L/N170E^ pull downs (Figure 2f). VPg - nCBP-2^W118L/N170E^ interaction was also observed through yeast two-hybrid assay (Figure 2g, h).

TuMV and CBSV belong to two different genera of *Potyviridae*, *Potyvirus* and *Ipomovirus*. It is possible that viruses from these distinct genera have evolved mechanistic differences in how their VPg proteins interact with host eIF4E-family *S* factors. To determine VPg protein conservation between potyviruses and ipomoviruses, VPg sequences from five major potyvirus clades were aligned with VPg sequences from seven of the eight known ipomovirus species (TomMMoV, SPMMV, SqVYV, CocMoV, CVYV, UCBSV, CBSV) (Figure S7). Across all VPg sequences analyzed, several highly conserved residues are observed, as are striking genus-specific differences; four stretches of amino acids are present only in one genus or the other (pink boxes). One of these stretches coincides with an α-α loop in PVY VPg thought to directly interact with the ventral side of eIF4E (Coutinho de Oliveira *et al*., 2019); in ipomoviruses this region is eight amino acids shorter. Additionally, a highly conserved aspartate residue (D77) necessary for TuMV VPg interaction with *Arabidopsis* eIF(iso)4E (Léonard *et al*., 2000), is not present in ipomoviruses, indicating this residue may have a less conserved role in interactions between ipomovirus VPgs and eIF4E-family proteins.

### A yeast two-hybrid screen identifies nCBP-2 mutants that lose VPg affinity

Since mutations from the known Arabidopsis resistance allele did not block nCBP-2 – VPg interaction, we performed a yeast two-hybrid screen for nCBP-2 mutants that lose affinity for VPg (Figure 3a). Mutant *nCBP-2* amplicons were generated by error-prone PCR and cloned into the bait vector by *in vivo* recombination in EGY48 yeast already containing both *lacZ* reporter and VPg prey vectors (Figure 3a). From roughly 1000 yeast transformants, 51 *nCBP-2* mutants were isolated that lost interaction with VPg (Table 2, Table S4). Forty-eight unique amino acid substitutions were observed across forty-six unique nCBP-2 mutants. Of the unique mutants, 27 had mutations affecting a single residue or the introduction of a frameshifting deletion while 19 contained two or three mutated residues. Nine residues were isolated more than once, with up to three unique amino acid substitutions identified. Mutants were then filtered to identify residues with varying degrees of conservation (Figure S8) across cassava eIF4E-family proteins resulting in 15 being re-cloned and validated for loss of VPg interaction in yeast using a quantitative β-galactosidase activity assay. All but two mutants, nCBP-2^L147H^ and nCBP-2^R196S^, were confirmed to confer total loss of β-galactosidase activity in yeast and suggest commensurate loss of VPg affinity (Figure 3b, 5b, S9). The R196S mutant yeast did not have any reduction of β-galactosidase activity relative to yeast expressing wild-type nCBP-2, while L147H mutant yeast exhibited an intermediate reduction in β-galactosidase activity. We further validated by western blot analysis that the yeast used for two-hybrid experiments expressed nCBP-2 variants and CBSV VPg (Figure S9).

**Fig. 3.**
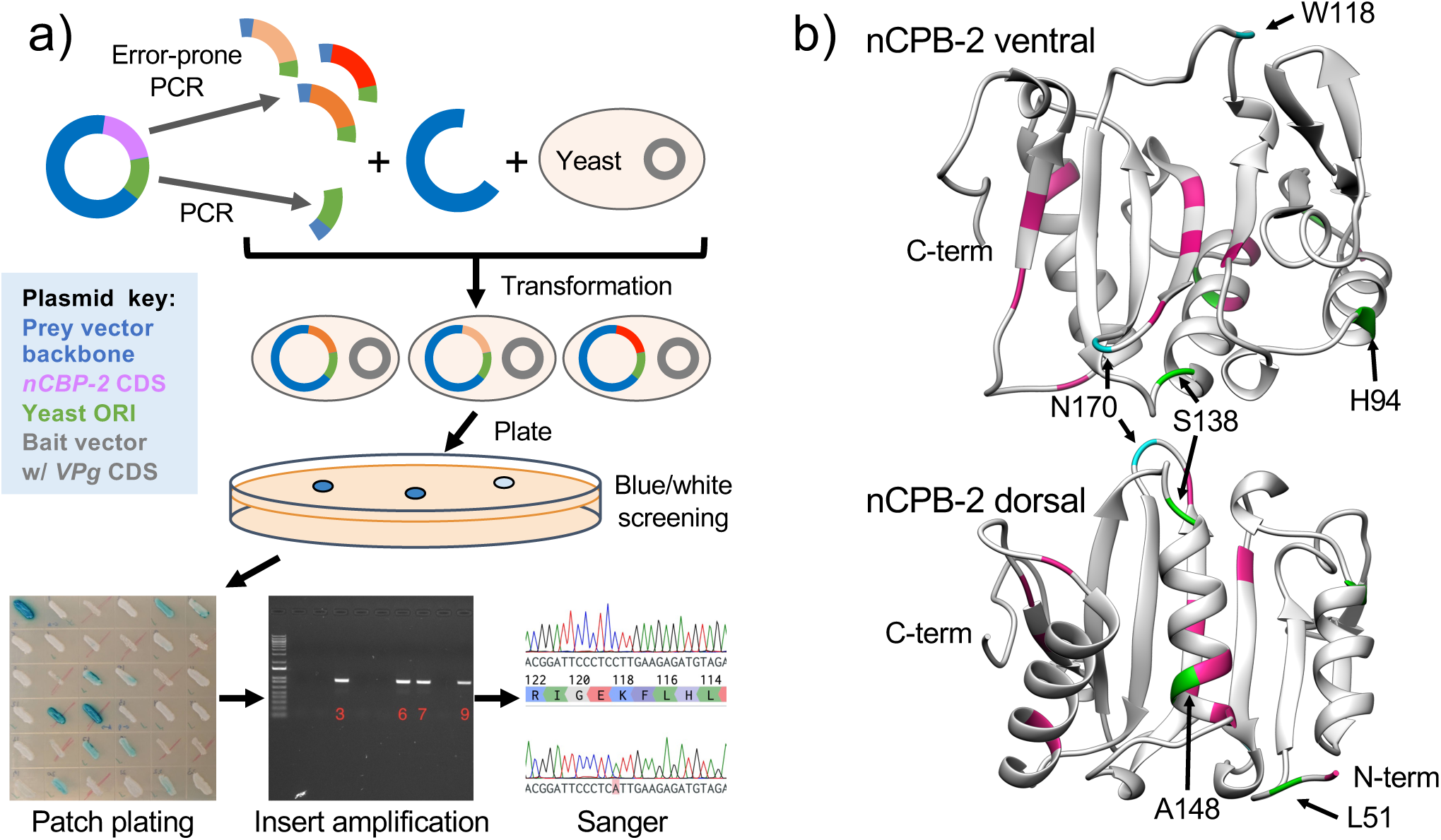
A yeast two-hybrid loss-of-affinity screen identifies nCBP-2 residues necessary for VPg interaction. a) Workflow for using a yeast two-hybrid approach to identify nCBP-2 mutants that lose affinity for CBSV Naliendele VPg. The resultant pEG202 prey vector encodes nCBP-2 fused to the LexA DNA binding domain. pJG4-5 encodes CBSV VPg fused to the B42 transcriptional activation domain. The EGY48 yeast strain used harbors the pSH18-34 reporter plasmid containing *lacZ* driven by the LexA operator. b) Ventral and dorsal views of cassava nCBP-2 structure produced using the Phyre2 server (Kelley *et al*., 2015). nCBP-2 structure of amino acids 49 through 226 was predicted using wheat eIF4E (PDB ID: 2IDR) as a template. Mutated amino acid residues in nCBP-2 mutants that lose VPg affinity and either rescue or fail to rescue *eif4e* yeast are highlighted in green and pink, respectively. W118 and N170, homologous with TuMV VPg interacting residues of *Arabidopsis* eIF(iso)4E, are highlighted in blue.

**Table 2.** Summary of nCBP-2 mutants found to lose affinity for CBSV VPg.

| No. loss-of-affinity mutants | Nucleotide substitutions per mutant (mean) | Nucleotide deletions per mutant (mean) | Unique alleles | No. unique AA substitutions | Frameshift mutants (unique/total) | Stop-gain mutants | Redundant AA substitution occurrences | AA residues with multiple occurrences of substitutions |
| --- | --- | --- | --- | --- | --- | --- | --- | --- |
| 51 | 1.37 | 0.15 | 46 | 48 <sup>a</sup> | 7/8 | 7 | 7 | 8 |
| <sup>a</sup> non-synonymous mutations downstream of frameshifting INDELs or stop-gain mutations are not counted |  |  |  |  |  |  |  |  |

### Translation initiation activity of isolated nCBP-2 mutants

Previous work with *Arabidopsis eif4e* or *eif(iso)4e* mutants suggested that both genes were functionally redundant for translation initiation. However, *eif4e* mutants exhibit subtle reductions in fertility and biomass in addition to delayed flowering time (Bastet *et al*., 2017, 2018). Similarly, simultaneous knockout of tomato *eIF4E1* and *eIF4E2* incurs a growth penalty (Gauffier *et al*., 2016). T93C is a yeast *eif4e* mutant with chromosomal *eIF4E* replaced with *LEU2* (Altmann *et al*., 1989). This yeast strain can survive on galactose media due to a plasmid-based *eIF4E* gene that is galactose inducible but is repressed in the presence of glucose (Figure 4a). Coding sequences of nCBP-2 mutants confirmed to lose affinity for CBSV VPg by quantitative yeast two-hybrid were cloned into pG1.1 under control of the constitutive yeast GPD promoter and transformed into T93C. Transfer of these T93C transformants from galactose- to glucose-containing media revealed that only three nCBP-2 mutants rescue loss of *eif4e*: nCBP-2^A20T/H94Q/L200P^, nCBP-2^L51F^, and nCBP-2^S138P^ (Figure 4bc, 5cd).

**Fig. 4.**
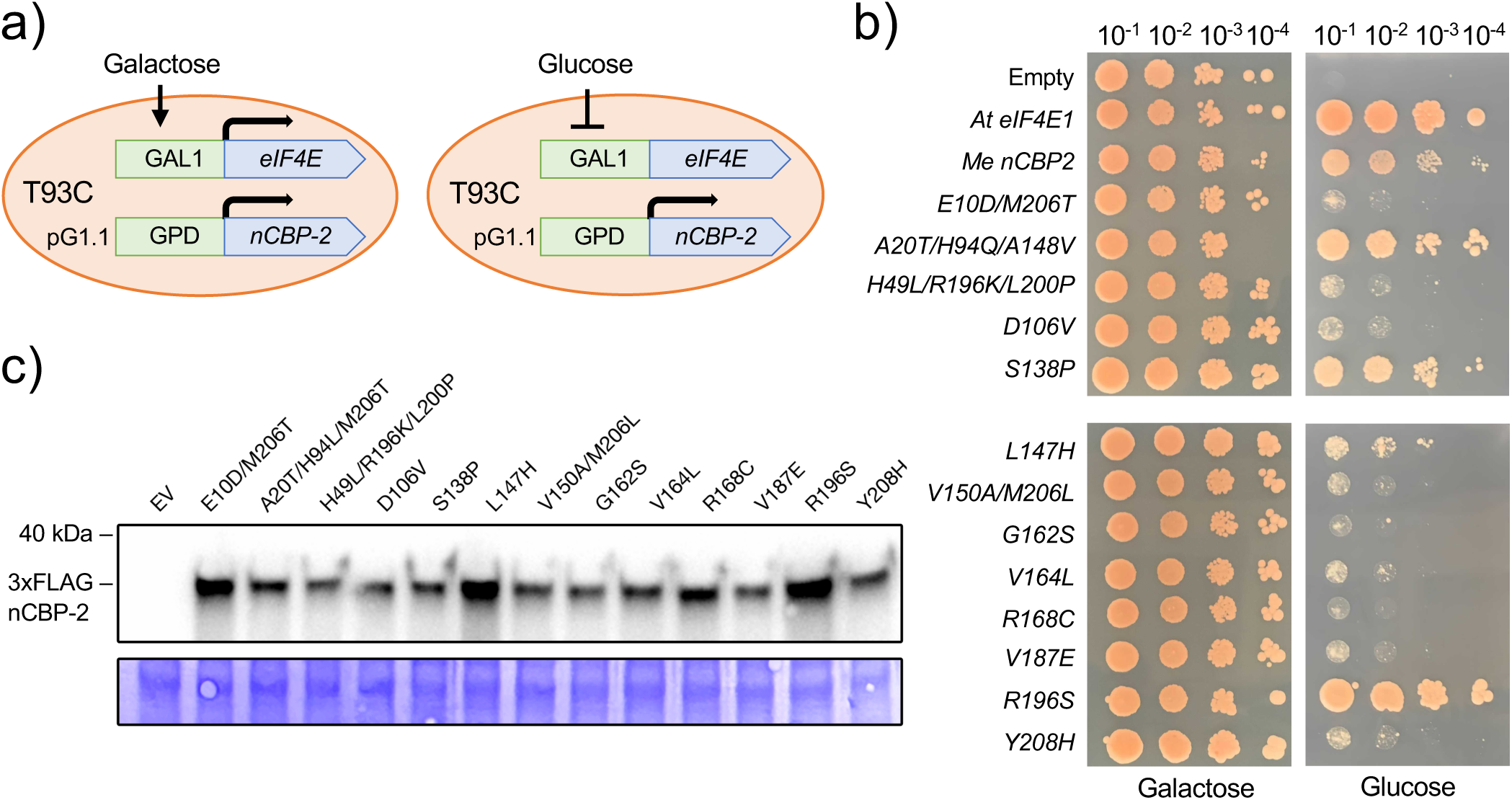
T93C *eif4e* yeast complementation by *nCBP-2* mutants. a) Schematic for testing translation initiation activity of *nCBP-2* variants expressed from the pG1.1 plasmid in T93C *eif4e* conditional mutant yeast. b) Left panels: T93C yeast transformed with pG1.1 empty vector or pG1.1 encoding various eIF4E-proteins grown on SD dropout media supplemented with galactose allowing for conditional expression of yeast eIF4E from the pGAL1-eIF4E plasmid. Right panels: same yeast strains from left panel grown on dropout media supplemented with glucose, preventing complementation by the pGAL1-eIF4E plasmid. c) Western blot analysis of 3xFLAG tagged nCBP-2 variant expression in yeast strains used in a). Coomassie stained membrane is presented for assessing loading control.

### Mutating the conserved HPL motif of nCBP-2 can affect VPg binding and disease

Among the three nCBP-2 mutants that retained translation initiation activity, nCBP-2^L51F^ stood out as having a mutation in the highly conserved HPL motif. In yeast, this motif participates in docking with eIF4G (Grüner *et al*., 2016, 2018). In addition, this amino acid is deleted in the nCBP-2^K45_L51del^ cassava mutant line that displayed increased resistance to CBSD (Figure 5a). The HPL motif is found in many plant eIF4E-family proteins and as such, a mutation of the leucine may be suitable for engineering resistance to other potyvirids. We tested the nCBP-2^K45_L51del^ mutant for interaction with VPg by quantitative yeast two-hybrid. Yeast expressing this mutant exhibited no β-galactosidase activity, like yeast expressing nCBP-2^L51F^ (Figure 5b, S10). We confirmed this implied loss of VPg interaction by observing strongly reduced VPg binding in GST pulldowns with both mutant proteins (Figure 6). However, comparing wild-type nCBP-2, nCBP-2^L51F^ and nCBP-2^K45_L51del^ for translation initiation activity in T93C yeast (Δ*eif4e*) showed that while nCBP-2^L51F^ was able to rescue loss of *eIF4E* in T93C it was to a lesser degree than wild type nCBP-2, and nCBP-2^K45_L51del^ was not able to complement T93C (Figure 5c, d).

**Fig. 5.**
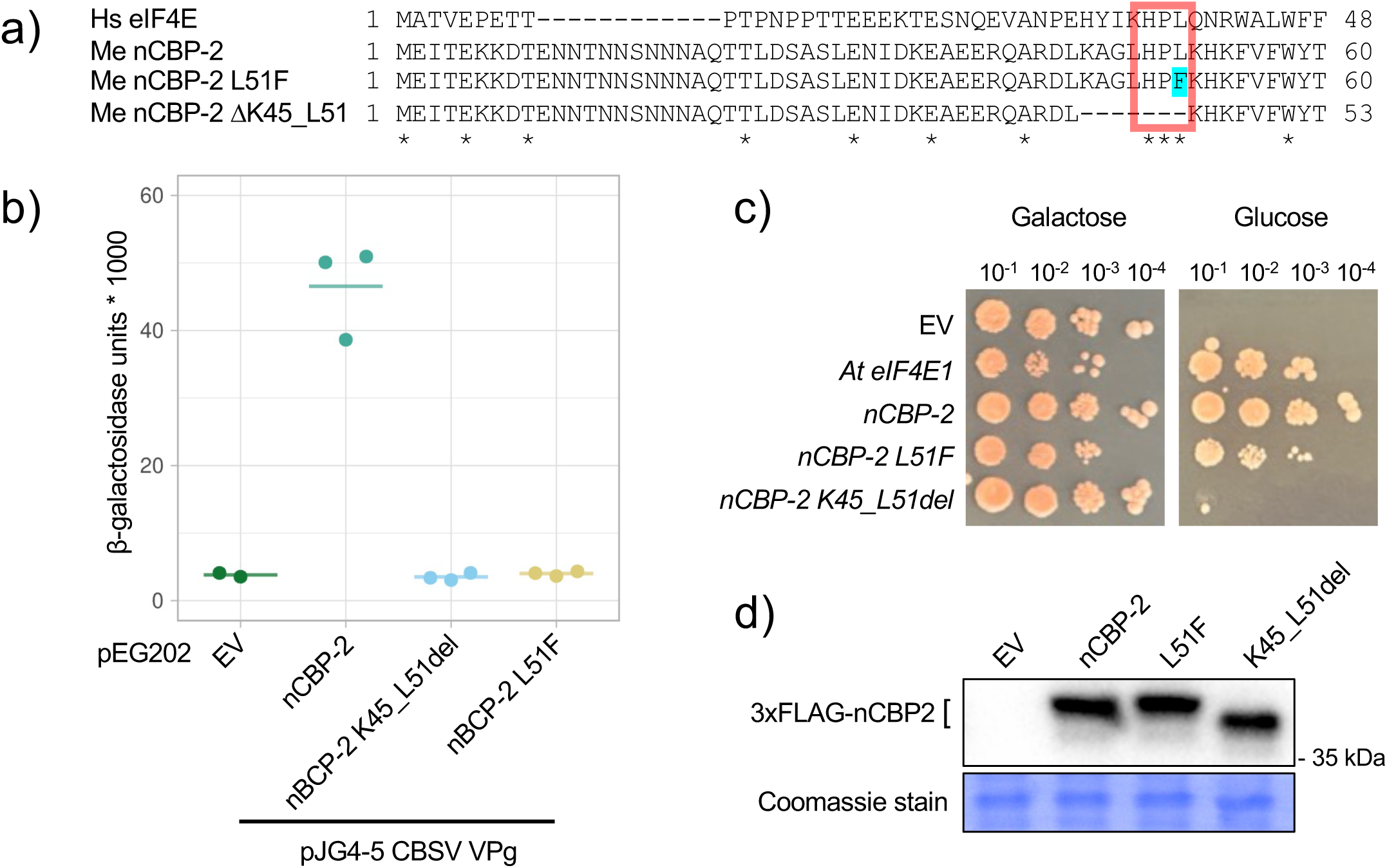
Effects of HPL motif mutation on nCBP-2 affinity for CBSV VPg and translation initiation. a) Alignment of human eIF4E protein sequence with variants of cassava nCBP-2. The conserved HPL motif is boxed in red and the L51F mutation in nCBP-2 is highlighted in cyan. Asterisks denote conserved amino acid residues. b) Quantitative yeast two-hybrid analysis of CBSV VPg interaction with nCBP-2 variants. c) T93C *eif4e* yeast transformed with pG1.1 encoding *Arabidopsis eIF4E1* or cassava *nCBP-2* variants under the control of a constitutive promoter. nCBP-2 variants are N-terminally tagged with 3xFLAG. The T93C yeast strain harbors a plasmid encoding galactose-inducible and glucose-repressible yeast *eIF4E*. d) Western blot analysis of nCBP-2 expression in yeast used in c). A segment of coomassie stained membrane is presented for loading control.

**Fig. 6.**
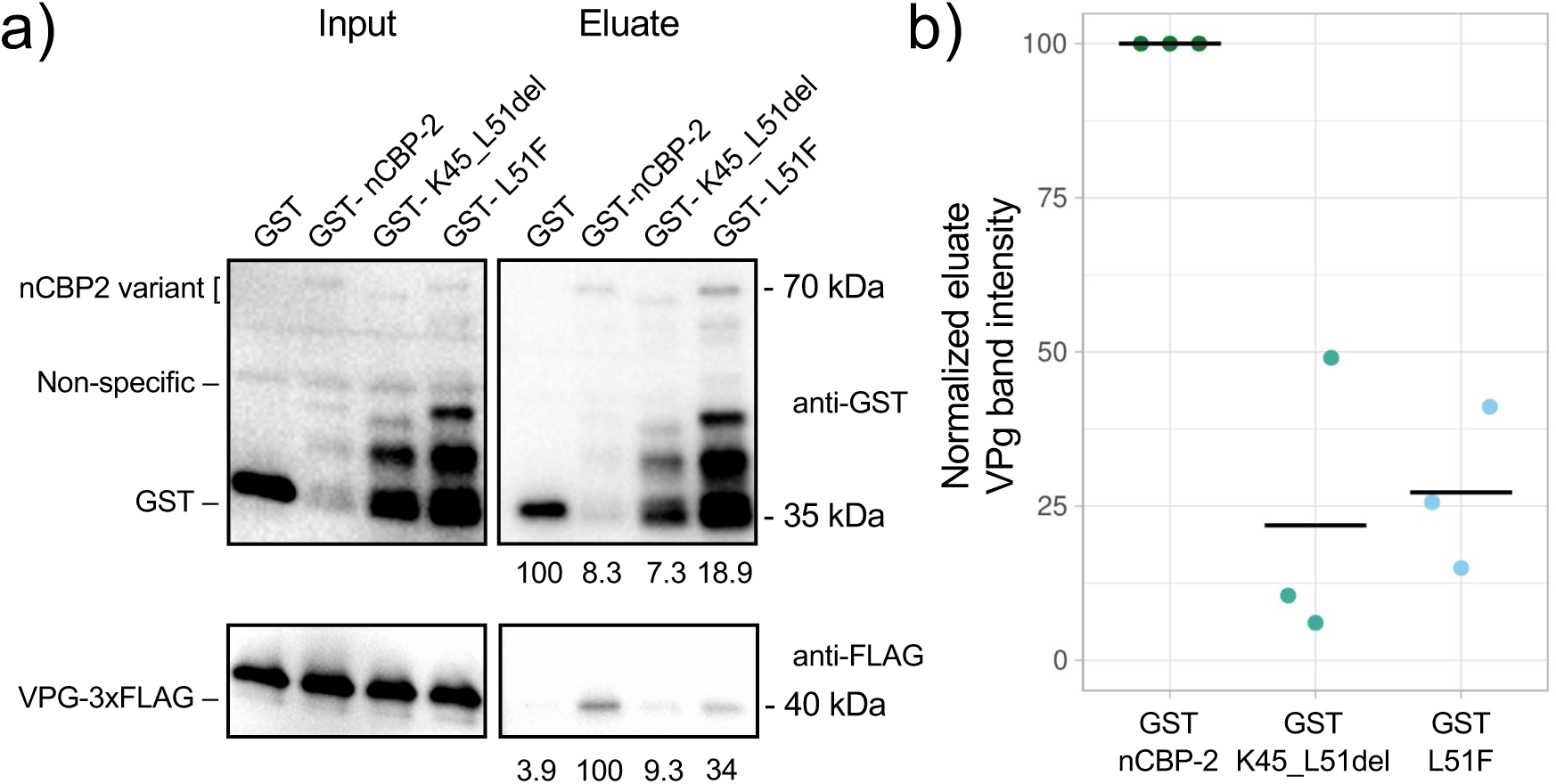
nCBP-2^K45_L51del^ and nCBP-2^L51F^ exhibit reduced affinity for CBSV VPg, *in vitro*. a) GST pulldown experiment assessing the ability of GST-nCBP-2 variants to bind CBSV VPg. Full-length protein eluate band intensities are presented underneath the eluate western blots and are expressed relative to the sample with the strongest band intensity. b) Analysis of eluate VPg band intensity across three GST-pulldown experiments. Band intensities are normalized to account for differing amounts of GST-fusion protein pulled down and expressed relative to VPg intensity in the GST-nCBP-2 sample.

### Mutating the HPL motif of other eIF4E-family proteins also disrupts VPg interaction

We also tested the effects of mutating the HPL motif on VPg interaction for the remaining cassava eIF4E-family proteins. Previously, we observed by yeast two-hybrid that CBSV VPg interaction with eIF(iso)4E-1 and eIF(iso)4E-2 is weak when compared to interaction with nCBPs (Gomez *et al*., 2019). Detecting these weak interactions requires incubating yeast for at least two days on solid growth media, in contrast to five hours of incubation in liquid media to detect interaction between nCBP-2 and VPg. All cassava eIF4E-family proteins and their HPL to HPF mutants were co-expressed in yeast on solid media for two days and then analyzed by quantitative β-galactosidase activity assay. nCBP-1^L46F^, eIF(iso)4E-1^L27F^, and eIF(iso)4E-2^L27F^ all exhibited reduced interaction with CBSV VPg compared to their wild type counterparts (Figure 7a, b). However, nCBP-1^L46F^ was found to not be expressed, while eIF(iso)4E-1^L27F^ consistently accumulated to lower levels relative to eIF(iso)4E-1 across experiments (Figure S11). eIF(iso)4E-2^L27F^ accumulated to similar levels relative to wild type protein. nCBP-1^L46F^ and eIF(iso)4E-2^L27F^ were then tested for their ability to rescue T93C *eif4e* yeast. eIF(iso)4E-2^L27F^ was able to rescue T93C, while nCBP-1^L46F^ could not, consistent with lack of nCBP-1^L46F^ accumulation in yeast two-hybrid experiments (Figure 7c).

**Fig. 7.**
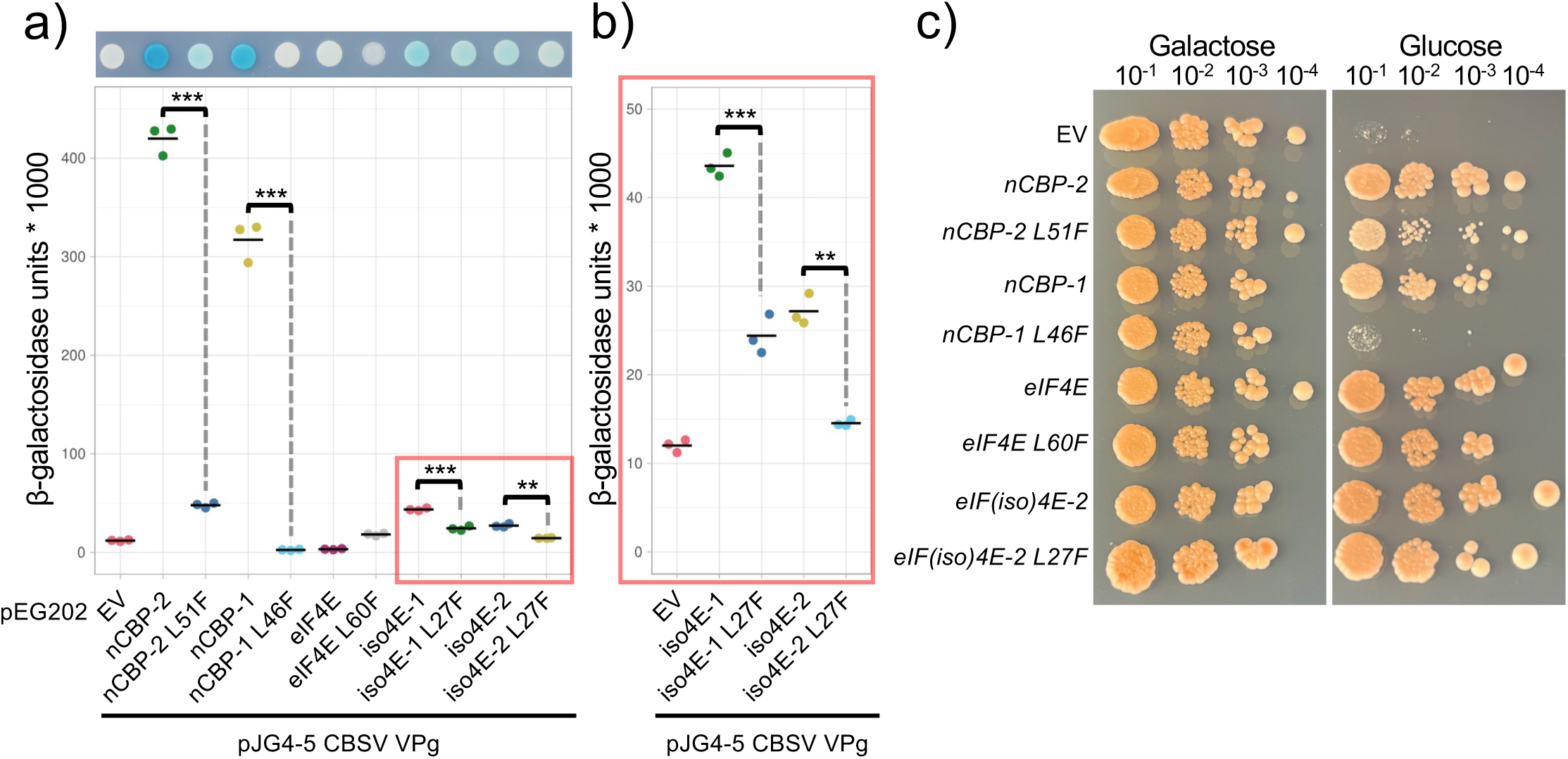
Effects of mutating the HPL motif leucine in cassava *eIF4E* and *eIF(iso)4E*. a, b) Quantitative yeast two-hybrid analysis of CBSV VPg interaction with wild type cassava eIF4E-famiy proteins alongside HPL to HPF mutants. Assays were performed after growing yeast under VPg inducing conditions for 48 hours. Results for eIF(iso)4E-1 and −2 interaction with CBSV VPg in panel a), highlighted with a red box, are enlarged in panel b). Statistical differences were detected using a Welch’s T-test with an alpha = 0.05. ** ≤ 0.01, *** ≤ 0.001. c) T93C *eif4e* yeast complementation assay with wild-type and mutant *eIF4E*-family genes from cassava

Leucine 51 of nCBP-2 corresponds to leucine 28 within *Arabidopsis* eIF(iso)4E. We hypothesized that recapitulating the L51F substitution in *Arabidopsis* eIF(iso)4E could have similar effects on TuMV VPg binding. When we tested this through quantitative yeast two-hybrid assays we observed that the L28F mutation strongly disrupted eIF(iso)4E interaction with TuMV VPg (Figure S12a). However, *Arabidopsis* eIF(iso)4E^L28F^ was not able to rescue T93C *eif4e* mutant yeast (Figure S12b, c).

## Discussion

*Potyviridae* is the largest plant-infecting family of RNA viruses comprised of 249 viruses across 12 genera (Walker *et al*., 2022). The genus *Potyvirus* is perhaps the best studied as it contains 206 of the 249 known viruses belonging to the family *Potyviridae*. In comparison, only eight species of viruses belonging to the genus *Ipomovirus* have been identified and much less is known about molecular ipomovirus-host interactions (Dombrovsky *et al*., 2014). While the genome structure and genes encoded by potyviruses and ipomoviruses are similar, and thus it is reasonable to expect a degree of conservation in virulence strategies, it is also likely that differences have arisen given the evolutionary divergence of the two genera. For CBSV in particular, we note that while eIF4E-family proteins are clear susceptibility factors, the breadth of family members that contribute to disease appears broader than typically found for potyviruses (Bastet *et al*., 2017). Furthermore, while resistance-conferring amino acid substitutions in eIF4E-family proteins have been effectively transferred between potyvirus-pathosystems, we find in this study that *Arabidopsis* eIF(iso)4E mutations that break TuMV VPg interaction fail to do so for the cassava nCBP-2 interaction with CBSV VPg. These observations highlight the need for further characterization of ipomovirus-host molecular interactions.

Structure-function analysis of VPg interaction with eIF4E-family susceptibility factors would strongly benefit from crystal or cryo-EM structures of the complex; these have yet to be successfully generated (Coutinho de Oliveira *et al*., 2019). Most recently, an NMR approach solved the structure of PVY VPg (Coutinho de Oliveira *et al*., 2019). The PVY VPg and human eIF4E interaction interface was also mapped through monitoring chemical shift perturbations of either protein during NMR spectroscopy. It is unclear how well these findings will translate to diverse potyvirid VPg interactions with cognate plant eIF4E-family proteins, especially for the plant specific eIF(iso)4E clade and considering significant divergence of ipomovirus VPg sequences from those of potyviruses. Thus far, attempts to design *eIF4E*-family resistance alleles have relied on inference-based approaches (German-Retana *et al*., 2008; Ashby *et al*., 2011; Kim *et al*., 2014; Bastet *et al*., 2018, 2019; Zafirov *et al*., 2021). In contrast, here we utilize a loss-of-affinity screen of randomly mutagenized nCBP-2 variants to identify key residues necessary for interaction with CBSV VPg. Only three nCBP-2 amino acid residues that we found to be necessary for interaction with VPg had previously been described in potyvirus-plant pathosystems. These amino acid residues, D106, V164, and R168 in nCBP-2, are highly conserved across eIF4E-family proteins. In pepper, the *pvr1^2^*allele of *eIF4E* is responsible for resistance to isolates of PVY and TEV, and the amino acid corresponding to nCBP-2 D106 is one of three causal amino acid substitutions (Kang *et al*., 2005). That aspartate in pvr1^2^ is substituted with asparagine (D to N). For cassava nCBP-2, two separate D106 substitutions were observed; D106V and D106H, the latter of which co-occurred with an A41T substitution. Only the nCBP-2^D106V^ mutant was tested for translation initiation activity and was found deficient in rescuing T93C *eif4e* yeast. It is unclear if a D106N mutation in nCBP-2, as found in pvr1^2^, would similarly break VPg interaction while preserving its translation initiation activity. nCBP-2 V164, where mutation to leucine disrupts CBSV VPg interaction, corresponds to V167 in eIF4E of pea. The pea eIF4E^V167A^ variant confers resistance to PSbMV and can rescue *eif4e* yeast. In contrast, cassava nCBP-2^V164L^ does not rescue T93C *eif4e* yeast. Whether the V164A mutation in nCBP-2 would have the same effects as in pea eIF4E is an open question. Lastly, for nCBP-2 R168, the previously described NMR structural study demonstrated that corresponding residues in human eIF4E and wheat eIF(iso)4E are involved in the interaction with *Potato virus Y* VPg (Coutinho de Oliveira *et al*., 2019). Like nCBP-2^D016V^ and nCBP-2^V164L^, nCBP-2^R168C^ also failed to rescue T93C.

Of the fourteen nCBP-2 loss-of-VPg-interaction mutants that we tested for translation initiation activity, eleven were nonfunctional. The three nCBP-2 variants that retained function were nCBP-2^A20T/H94Q/A148V^, nCBP-2^L51F^, and nCBP-2^S138P^. For nCBP-2 ^A20T/H94Q/A148V^, A20 resides outside of the conserved core of eIF4E family proteins while H94 and A148 are present in alpha helices found on the dorsal side of nCBP-2; further work is required to identify which individual or combination of these amino acids are required for CBSV VPg interaction. S138 is similarly located in a dorsal alpha helix while L51 is closer to the dorsal side and located N-terminal of a beta strand. Evidence from a study on PVY VPg interaction with human eIF4E indicates that VPg interfaces with the ventral side of eIF4E (Coutinho de Oliveira *et al*., 2019). If this also applies to CBSV VPg interaction with nCBP-2, the four aforementioned amino acid residues likely affect VPg binding in an indirect manner. Of particular importance for this study, the L51F substitution impacts the HPL motif, which was also affected in our cassava *ncbp-1 nCBP-2^K45_L51del^* line. This led us to test the importance of that motif for CBSD symptom severity. Unlike nCBP-2^L51F^, nCBP-2^K45_L51del^ did not rescue T93C *eif4e* deficient yeast. Surprisingly, despite the inability to strongly interact with VPg and complement *eif4e* activity, the *ncbp-1 nCBP-2^K45_L51del^* root disease phenotype was intermediate to wild type and *ncbp-1 ncbp-2*, indicating that this mutant version of *nCBP-2* is sufficient for the development of some disease symptoms. A similar result has also been observed in pea *eIF4E* mutants where of eight mutants unable to rescue *eif4e* mutant yeast, two were still able to confer susceptibility to PsbMV (Ashby *et al*., 2011). In such mutants, it may be that low levels of residual translation initiation activity are sufficient for completion of the viral life cycle. Lastly, we note that in a more sensitive yeast two-hybrid assay, while VPg interaction with nCBP-2^L51F^ resulted in eight-fold less reporter activity than with nCBP-2, the strength of this interaction appeared similar to that with eIF(iso)4E-1. This suggests that mutating additional nCBP-2 residues may be necessary for total elimination of VPg binding and for optimal resistance *in planta*.

In cassava, the eIF4E-family has five members, and we previously demonstrated that all five proteins interact with CBSV VPg by co-immunoprecipitation (Gomez *et al*., 2019). We now find, through pulldown experiments with purified proteins, that nCBP-1, nCBP-2, and eIF4E can directly interact with CBSV VPg. This validates our approach for identifying nCBP-2 residues crucial for VPg interaction. In contrast, our pulldown experiments with VPg and eIF(iso)4E-1 and −2 yielded inconsistent results across experimental replicates. This is difficult to interpret, though it is possible that levels of recombinant protein activity vary between purified batches. Further optimization of purification conditions for cassava eIF(iso)4E-1 and −2 may resolve this issue. Regardless, our *in vitro* data lay the groundwork for a more granular investigation of differences in cassava eIF4E-family protein affinity for CBSV VPg; this may be achieved using techniques like surface plasmon resonance or isothermal titration calorimetry (Upadhyay et al., 2024).

To determine the contribution of each *eIF4E-*family gene to CBSV symptoms, and therefore the biological relevance of interaction with VPg, we have used CRISPR/Cas9 to generate single mutants of all five genes, as well as some double mutants, in cassava variety TME419. Notably, the *ncbp* double mutant has near total reduction of storage root necrosis, commensurate with strong reductions in viral titer. Of the two *nCBP* genes, *nCBP-2* appears to be the main driver for the improved disease response as evidenced by the results of the single *nCBP* mutants, and that *nCBP-2* is more highly expressed than *nCBP-1* (Gomez *et al*., 2019). As noted in our previous work, the expression difference between *nCBP-2* and *nCBP-1* is even more pronounced in storage roots and may be the reason for differential disease response in roots and aerial tissues of *ncbp-2* and *ncbp-1 ncbp-2* mutants (Gomez *et al*., 2019). The disease data are consistent between mutants in both 60444 (Gomez *et al*., 2019) and TME419. Beyond the *nCBP* clade, the *eIF(iso)4E* clade appears to have a small contribution to disease as aerial symptom severity and storage root necrosis is slightly reduced when both *eIF(iso)4E-1* and *eIF(iso)4E-2* are simultaneously knocked out. This finding also argues against the *ncbp-1 ncbp-2* root disease phenotype being simply due to additively knocking out family members. It remains to be seen if higher order mutant combinations would have an even greater effect on reducing CBSD disease severity. It will also be important to test the utility of *ncbp-1 ncbp-2* mediated resistance against the natural diversity of cassava brown streak viruses in field trials.

A key consideration going forward is that any benefits conferred by mutations in eIF4E-family proteins with respect to viral resistance may also be accompanied by growth and developmental penalties (Gauffier *et al*., 2016, Bastet *et al*., 2017). Of the four residues identified in *nCBP-2* mutants that may avoid this penalty, three (L51, H94, and A148) are conserved within all cassava eIF4E-family proteins, and the fourth (S138) is conserved within three members. Targeted base editing of some or all of these residues across the entire eIF4E-family may allow for improved CBSV response while maintaining essential protein translation functions. Indeed, we demonstrate here that transferring the L51F mutation from *nCBP-2* into cassava and *Arabidopsis* eIF(iso)4E proteins results in loss of affinity for corresponding VPgs. As base editing approaches mature alongside improvements with gene targeting, combining these approaches with loss-of-VPg interaction screens may facilitate isolation and implementation of novel *eIF4E-*family resistance alleles.

## Materials and Methods

### Plasmid vector construction

gRNAs for CRISPR/Cas9-based mutagenesis of cassava *eIF4E-*family genes were selected using the CRISPOR webtool (Concordet & Haeussler, 2018). Cas-OFFinder was then used to screen gRNAs for off-target activity against a TME419 reference based genome assembly produced using data (SRX1393295) from Bredeson et al. and the AM560-2 v6.0 reference genome (Table S1) (Bae *et al*., 2014; Bredeson *et al*., 2016). gRNAs with off-targets containing less than 2-mismatches were excluded from consideration. Finally, gRNAs were cloned into the pDIRECT 21C CRISPR binary vector as previously described by (Čermák *et al*., 2017).

### Plant transformation, genotyping, and growth parameters

Cassava cultivar TME419 was transformed as previously described (Veley *et al*., 2023). DNA was extracted from recovered transformants using the CTAB procedure. PCR was then used to amplify 240 to 428 bp regions encompassing the gRNA binding sites. Purified amplicons were sent to GENEWIZ for amplicon sequencing. Amplicon sequencing data was analyzed using the CRIS.py python program (Connelly & Pruett-Miller, 2019). Plantlets of transgenic lines were then maintained in tissue culture (Taylor *et al*., 2012).

### Viral challenge and disease phenotyping

Cassava disease challenges with CBSV isolate Naliendele, including inoculation, aerial and below ground disease severity assessments, and qPCR-based quantification of storage root viral titer, were carried out as described previously (Gomez *et al*., 2019). For data analysis, wild type disease severity was compared across all challenges and no significant differences were found (Figure S1). Data for all challenges were then aggregated to increase sample sizes for statistical analysis.

### Co-immunoprecipitation and GST-pulldown experiments

Co-immunoprecipitation was performed as described previously (Gomez *et al*., 2019). For GST-pulldown experiments, cassava *eIF4E-*family genes in pENTR/D-TOPO, from Gomez et al., were cloned into pDEST15 (Invitrogen) by Gateway recombination (Invitrogen). pDEST15 encoding cassava *eIF4E-*family genes and pGEX-4T-1 (encoding unfused GST, Cytiva) were transformed into BL21-AI E. coli (Invitrogen) and transformants were used to start overnight seed cultures. Seed cultures were then used to inoculate 250 mL of LB broth. Once inoculated cultures were grown to an OD600 of 0.4, IPTG (pGEX-4T-1) or arabinose (pDEST15) were added to final concentrations of 0.5 mM or 0.2%, respectively, to induce recombinant protein production. Induced cells were incubated for four hours at 28°C. Cells were then pelleted and resuspended in 4 mL of lysis buffer (125 mM Tris, 150 mM NaCl, 1 mM DTT, 5 mM MgCl_2_, 10 mM CaCl_2_, pH 7.4) supplemented with 1X cOmplete protease inhibitor (EDTA-free, Roche). Lysozyme (GoldBio) was added to a concentration of 1 mg/mL and the mixture was allowed to incubate at room temperature for 15 minutes. 50 uL DNAse I (Thermo) was then added to the mixture, followed by sonication on ice to ensure complete lysis and elimination of DNA induced viscosity. The resulting cell lysate was centrifuged at 4°C, maximum speed, for 20 minutes. 3.5 mL of clarified lysate was transferred to a new tube and EDTA was added to a final concentration of 1 mM. GST and GST-eIF4E proteins were then purified from clarified lysate using glutathione magnetic agarose beads (Pierce) according to manufacturer’s protocol. Glutathione was removed from GST and GST-fusion protein eluates by buffer exchange with Amicon Ultra centrifugal filters (Millipore). CBSV 6xHis-VPg-3xFLAG-6xHIS was produced and purified for pulldown experiments as described in (Gomez *et al*., 2019). GST pulldown was performed by mixing 3 ug of GST or GST-eIF4E proteins with 3 ug of VPg and 10 uL of resuspended glutathione magnetic agarose beads (Pierce) in buffer A (125 mM Tris, 150 mM NaCl, 10 mM DTT, 1 mM EDTA, pH 7.4) supplemented with 1X cOmplete protease inhibitor (EDTA-free, Roche). Samples were incubated at 4°C for one hour. Beads were then washed four times with buffer A and proteins were eluted with Laemmli sample buffer for analysis by western blotting. Antibodies used in this study are described in Table S2.

For comparison of VPg pulldown by nCBP-2 variants across experiments, nCBP-2 and VPg eluate band intensities were first measured using Image Lab (Bio-Rad). Within an experiment, eluate VPg band intensities were first adjusted to correct for differing pulldown efficiencies of each nCBP-2 variant using the following equation where VPg_X_ is VPg band intensity for sample X and GST_X_ is full length GST fusion protein band intensity for sample X:

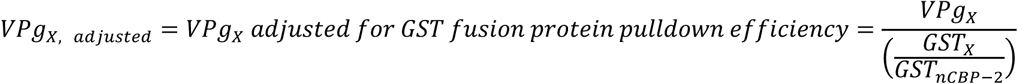

Adjusted VPg intensities were then expressed relative to the VPg intensity of the wild-type nCBP-2 sample for that experiment:

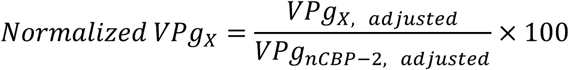

### Yeast two-hybrid assays

All yeast two-hybrid experiments were performed using the EGY48 strain harboring the pSH18-34 *lacZ* reporter plasmid along with pJG4-5 and pEG202 bait and prey plasmids (Golemis & Khazak, 1997). pJG4-5 *CBSV VPg* bait plasmid and pEG202 *nCBP-2* prey plasmids were created by using In-Fusion cloning (Takara) to insert CBSV VPg or nCBP-2 coding sequences into pJG4-5 or pEG202 linearized with EcoRI and XhoI. For qualitative yeast two-hybrid experiments, EGY48 pSH18-34 yeast transformed with bait and prey plasmids were first selected for on SD –ura –trp –his media supplemented with 2% glucose (Gietz & Schiestl, 2007). Transformants were then plated onto SD –ura –trp –his media supplemented with 2% galactose, 1% raffinose, 1X BU salts, and X-gal. Blue/white screening was performed three to four days later. Quantitative yeast two-hybrid was performed using a spectrophotometric β-galactosidase activity assay (Amberg *et al*., 2006). To extract protein for western blotting, 1 mL of yeast at OD600=2 was pelleted, resuspended in 0.1M NaOH, incubated at room temperature for five minutes, pelleted, resuspended in Laemmli sample buffer, incubated at 95C for five minutes, and pelleted again. The resulting supernatant was used for SDS-PAGE and western blot analysis.

To screen for nCBP-2 mutants that lose affinity for CBSV VPg, pEG202 *nCBP-2* was used as template for mutagenic PCR (Genemorph II, Agilent). Primers immediately flanking the second and penultimate codon of *nCBP-2*, preserving the start and stop codons during mutagenic PCR, were used with the low mutation rate protocol from the Genmorph II kit (Agilent). pEG202 empty vector was then digested with EcoRI and SacI and the fragment containing the yeast *HIS* marker was gel purified (Qiagen). A third pEG202 fragment containing the yeast 2*μ* origin of replication was produced by PCR with ends that overlap the *nCBP-2* mutant amplicons and pEG202 *HIS* marker fragment by 30 base pairs; co-transformation of all three fragments into yeast result in assembly of an intact pEG202 *nCBP-2* plasmid through yeast recombinational cloning. Mutant *nCBP-2* amplicons, pEG202 *HIS* fragment, and pEG202 2*μ* fragment were mixed in a 4:1:1 molar ratio and co-transformed into EGY48 yeast harboring the pSH18-34 and pJG4-5 *CBSV VPg* plasmids. Transformed yeast were directly plated onto SD –ura –trp –his media supplemented with 2% galactose, 1% raffinose, 1X BU salts, and X-gal. Four days post transformation, white or light blue colonies were restreaked onto a new plate for further visual assessment of reduced β-galactosidase activity. The yeast were then checked by PCR for the presence of full length *nCBP-2* insert in pEG202 using vector backbone specific primers. *nCBP-2* mutant alleles were then amplified from insert positive yeast using Q5 Hot Start polymerase (NEB) and analyzed by Sanger sequencing.

### Yeast *eif4e* complementation assay

3xFLAG and nCBP-2 or mutant nCBP-2 coding sequences were cloned into pG1.1 linearized with NcoI and BamHI using In-Fusion (Takara) to produce pG1.1 *3xFLAG-nCBP-2*. *3xFLAG* was amplified from pB7-HFC (Huang *et al*., 2016) while *nCBP-2* variants were amplified from pEG202 yeast two-hybrid plasmids. pG1.1 plasmids were then transformed into the conditional yeast *eif4e* mutant strain T93C (*eif4e*::LEU2 *ura3 trp1 leu2* [pGAL1-eIF4E *URA3*]) (Altmann *et al*., 1989). Transformants were plated on SD –ura –trp dropout media containing 2% galactose and 1% raffinose and allowed to grow for five to seven days. Colonies of transformed yeast were then resuspended in sterile water and a ten-fold dilution series ranging from OD600 = 10^-1^ through 10^-4^ was plated on SD –ura –trp dropout media containing 2% glucose to examine for rescue by pG1.1 encoded *eIF4E-*family gene variants. Plates were imaged after 5-7 days.

### SDS-PAGE and western blotting

Proteins in Laemmli buffer were run on TruPAGE Triethanolamine-tricine (SIGMA, discontinued) or Novex WedgeWell Tris-glycine (Thermo Fisher) SDS-PAGE gels according to manufacturer’s instructions. Samples were run alongside Benchmark Pre-stained or PageRuler Prestained protein ladders (Thermo Scientific). We note that kDa markers were observed to shift up to 15 kDa depending on gel and buffer chemistry. Proteins were then transferred onto Immobilon-PSQ PVDF membranes (Sigma) at 100V constant for one hour. After blocking in 5% milk in TBST, proteins were detected by probing with antibodies listed in Table S2. After imaging, membranes were immediately rinsed in TBST and stained with Coomassie R250 to check for equal loading.

## Acknowledgements

We would like to thank the DDPSC PGF team for their hard work in caring for our plants. We thank Claire Albin for her assistance with CBSV challenges and Nigel Taylor for allowing us to use Taylor Lab greenhouse space. Yeast strain T93C, pG1.1 At_eIF4E1, and pG1.1 empty vector were generously provided by Dr. Karen S. Browning. Z.D.L. would like to thank Susan Lin, Wei-Yang Lin, Chun-Hwa Lin, and Jocelyn Lin for their love and support. Z.D.L. was funded by USDA NIFA award 2019-67012-29626 and the Bill & Melinda Gates Foundation (OPP1125410, R.B.S., S.E.J., J.C.C.). E.J.D., Z.V.B., H.T., E.H., and A.M. were funded by the DDPSC NSF-REU program, DBI-1659812, −2050394, −1659812, and −2244275.

## Author Contributions

Conceptualization: Z.D.L.; Methodology: Z.D.L., J.C.C., R.S.B.; Investigation: Z.D.L., M.K.S., G.L.H., E.D.M., Z.V.B., K.B., H.T., E.H., G.J., K.B.G.; Visualization: Z.D.L.; Writing – original draft: Z.D.L.; Writing – reviewing & editing: Z.D.L., K.B.G., J.C.C., R.S.B.

## Data Availability

The source data for all results presented in this study are provided in accompanying supplementary files.

