## Supplemental Figures for "Targeted and random mutagenesis of cassava brown streak disease susceptibility factors reveal molecular determinants of disease severity"

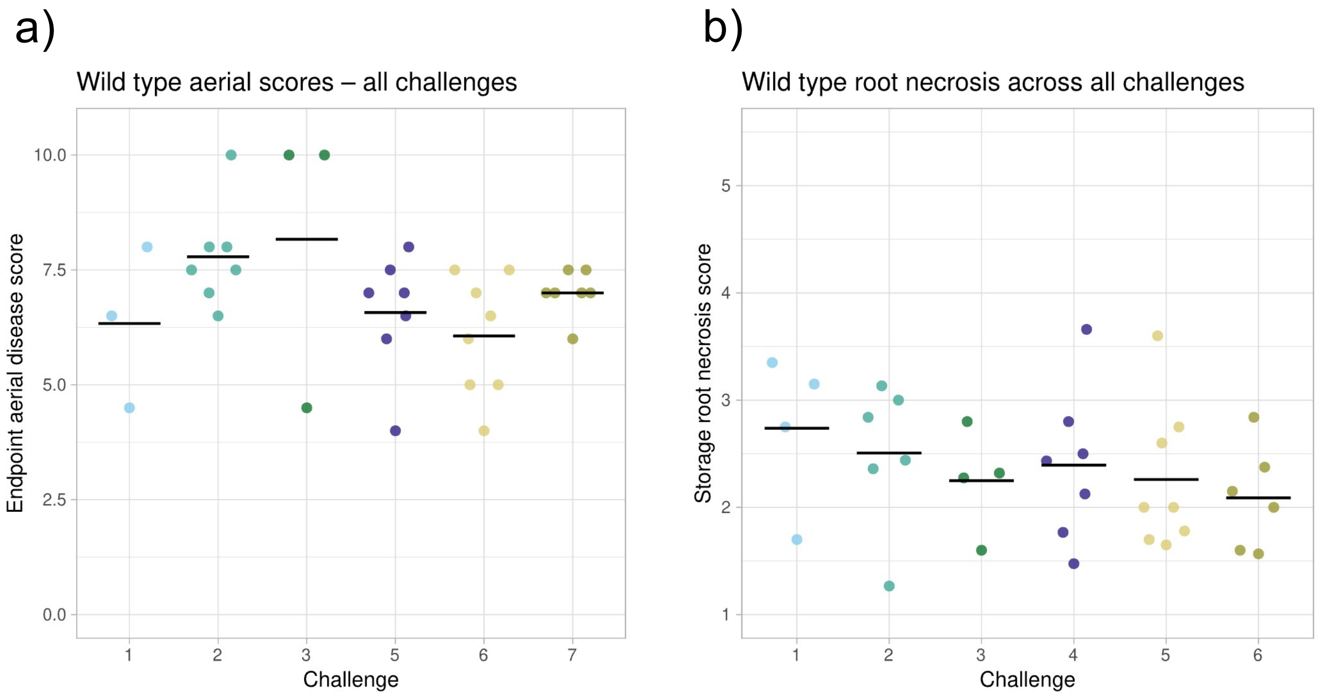


Fig. S1 Wild type aerial and storage root disease scores across all challenges
a, b) Aerial and storage root disease severity scores for wild type TME419 across all challenges. No statistically significant differences between challenges were detected by the Kruskal-Wallis test.

*eIF4E-1* exon 1

ATGGCGGCGGAAGAGCCATTGAAATCAACTACTGAAGAAACACCGAATCCAAACCTTAACTCTAATCCTAGGGCACTAGATGACGTTAATGATGATGAGCCGGAAGAAGGAGAGATCGTCGGAGATGAGGAATCATCCGCCAAGAAATCCAGCGCTGTAACC
TACCAGCCGCACCCTCTCGAGCATCAGTGGACGTTCTGGTTCGATAACCCTACTGCTAAGTCTAAGCAAGCCACCTGGGGAAGCTCTATGCGTTCTATCTATACTTTTGCTACTGTTGAGGAGTTTTGGAG

*eIF(iso)4E-1* exon 1

ATGGCAACCGAAACAGCAACAGAAGGAAGCGCCACGGAGGCCACAGCTACCGGTGTTGAGAAGCCGCTGCAGCACAAGCTAGAGAGGAAATGGACCTTCTGGTTCGATAATCAATCCAAGCCCAAACAAGGCGCCGCCTGGGGCAGCTCTCTCCGCAAGGT
CTACACCTTTGACACCGTTGAAGAATTTTGGTG

*eIF(iso)4E-2* exon 1

ATGGCAAGCGAAACCGCAATAGAAGGCACCGCGACAGAGGCTACAGCTTCTGGCGCAGAGCAGCCGCAACAGCACAAGCTAGAGAGGAAGTGGACGTTCTGGTTTGATAATCAATCCAAGCCTAAACAAGGTGCCGCCTGGGGCACCTCTCTCCGCAAGAT
CTACACGTTCGATACTGTCGAAGAATTTTGGTG

*nCBP-1* exon 1

ATGGAGATCACAGAGGAGGAAACAGAAAACAATAACTATAGCAATAATAATGCTCGAACTGCATCATCACCGGATAATATCGATAAAGTAGCCGAAGAACGCCAAGCAAGGGACCTCGAAGCTGGTTTGCATCCTCTCAAG

*nCBP-2* exon 1

ATGGAGATCACTGAGAAGAAGGATACAGAGAACAACACAAACAACAGCAATAATAATGCTCAAACGACATTGGATTCAGCATCACTTGAGAACATAGACAAAGAAGCCGAAGAACGCCAAGCACGCGACCTCAAAGCTGGGTTGCATCCTCTCAAG

Figure S2. gRNAs used in this study

gRNA target sequences are highlighted in yellow or green on the exon 1 sense sequence of cassava *eIF4E*-family genes. PAM sequences are highlighted in blue. The *nCBP-2* gRNAs highlighted in yellow and green were used to generate the *ncbp-1/ncbp-2* and *ncbp-1/nCBP-2*$^{K45\_L51del}$ mutants, respectively.

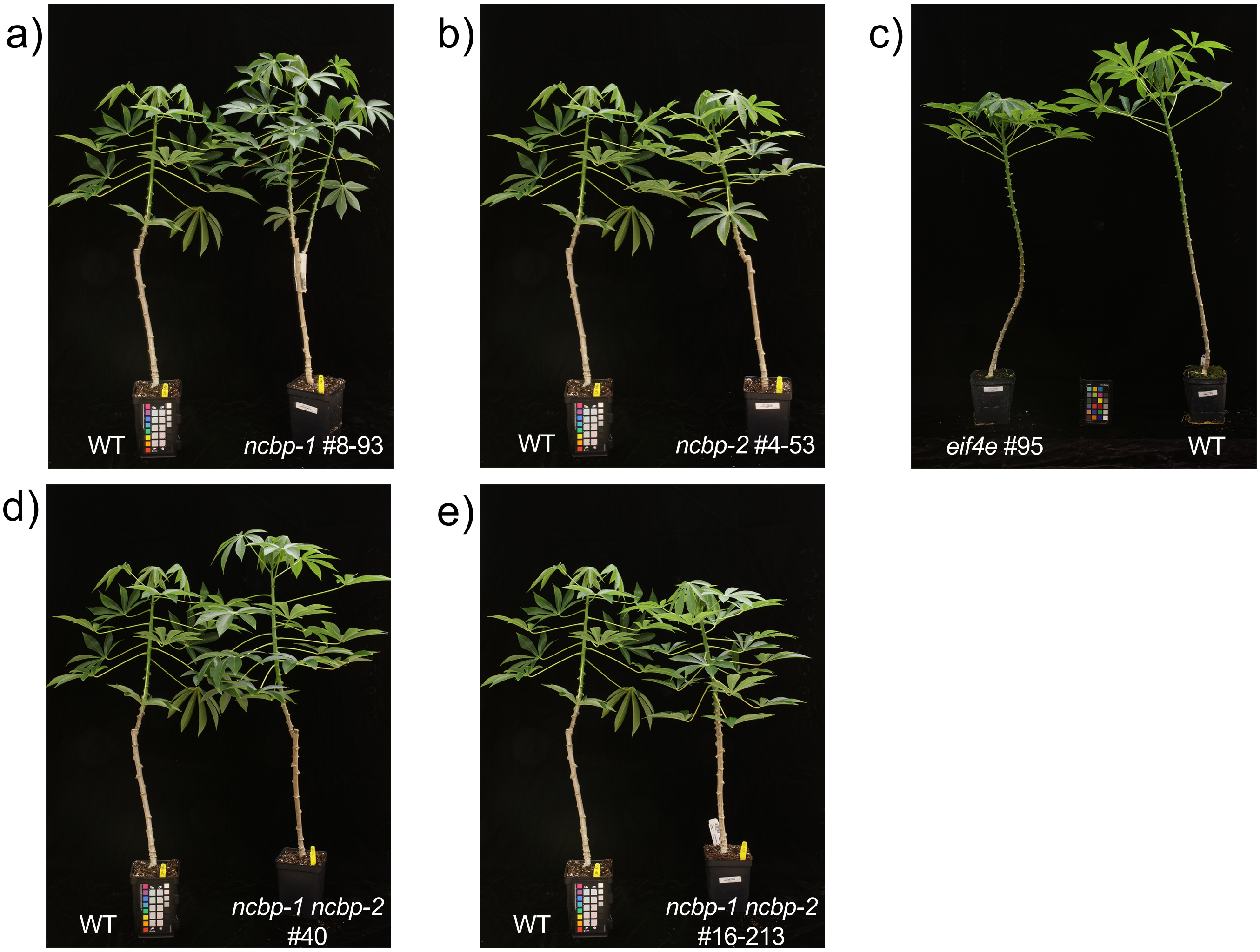


Fig. S3 *eIF4E*-family mutant cassava plants

a-e) Photos of of representative *eIF4E*-family mutants in TME419 background 6 months most transfer to soil

*eif(iso)4e-1 eif(iso)4e-2*
#172

WT

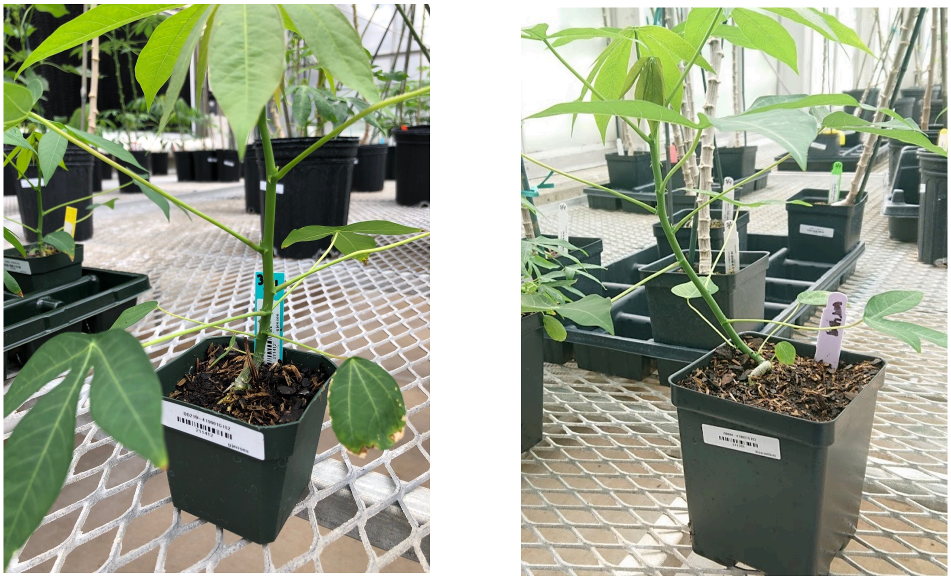

Fig. S4 Representative *eif(iso)4e-1 eif(iso)4e-2* double mutant and wild type TME419 six weeks post transfer to soil

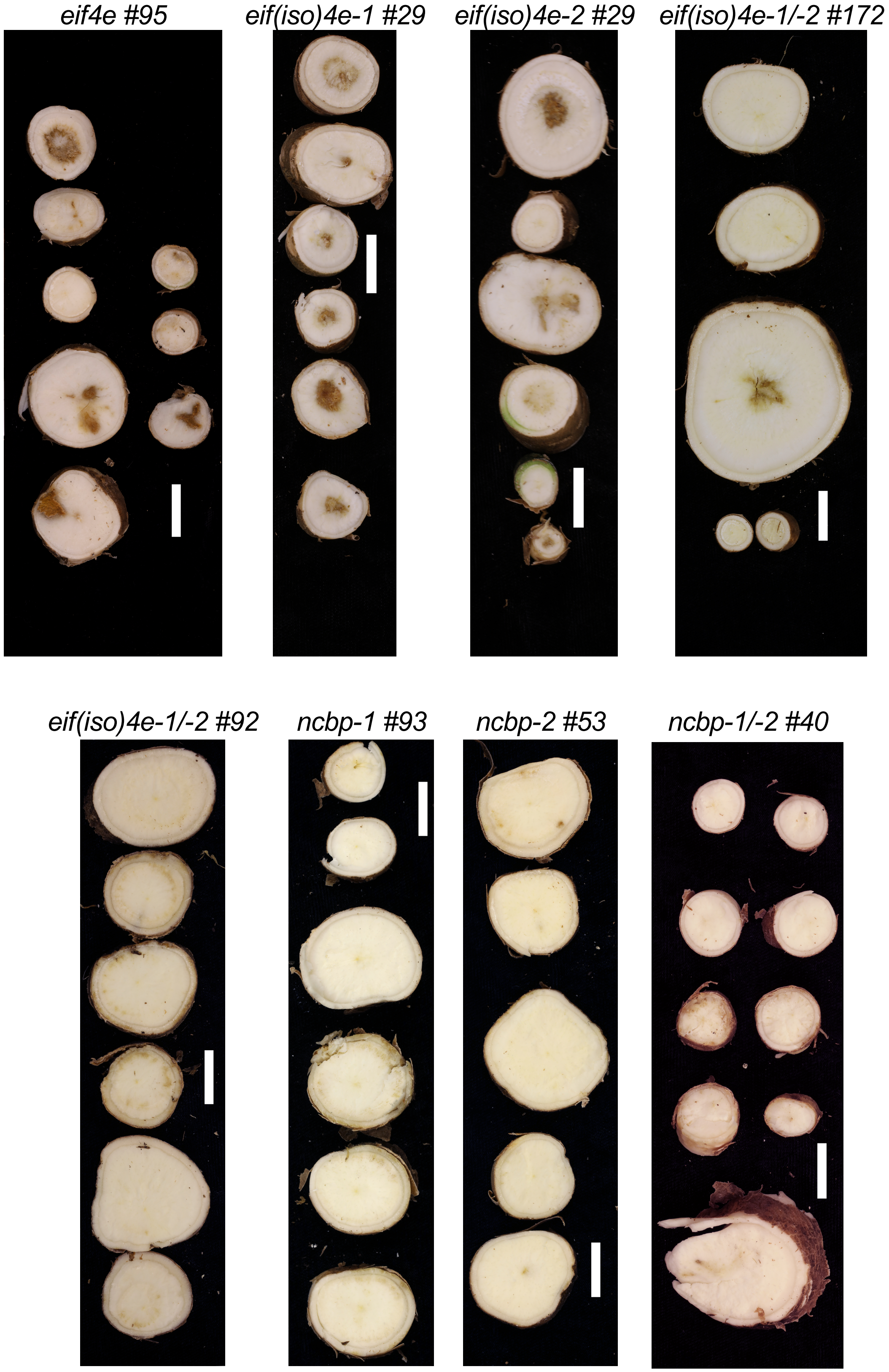


Figure S5. Storage root photos of CBSV infected mutant cassava

Storage root sections from CBSV infected plants. Each panel is from a single plant and is representative of median observed disease severity for each cassava *eIF4E*-family mutant line. White scale bar denotes 1.3 cm.

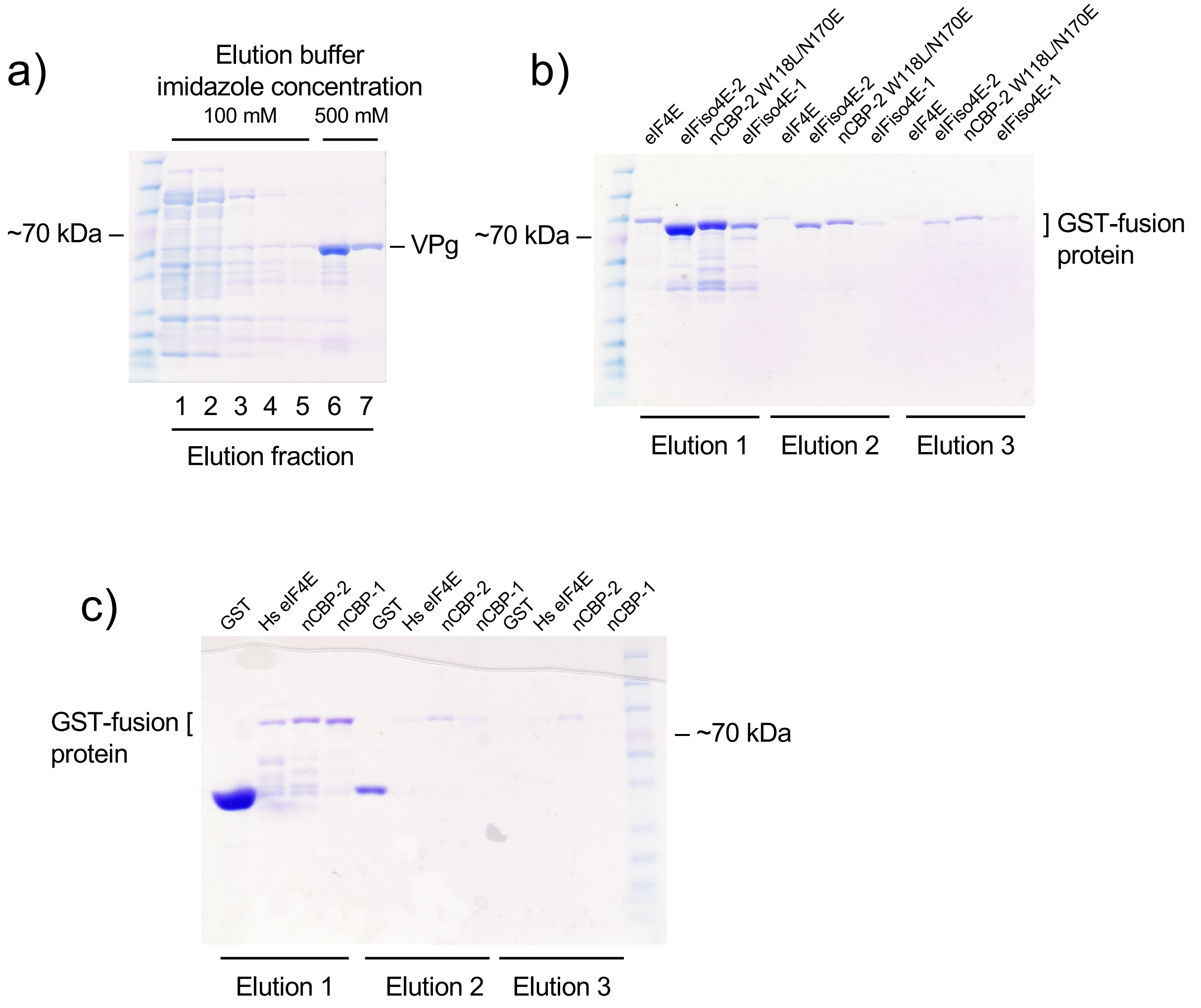


Figure S6. Purified proteins for GST pulldown assays

a-c) SDS-PAGE gels run with various elution fractions of purified proteins used in GST pulldown assays. 3xFLAG and dual-6xHis tagged CBSV Naliendele VPg was purified on Ni-NTA agarose and eluted with five sequential washes of 100 mM imidazole elution buffer followed by two sequential washes of 500 mM imidazole elution buffer. GST and GST-fusion proteins were purified with magnetic glutathione conjugated beads and eluted with three sequential washes of 50 mM glutathione elution buffer.

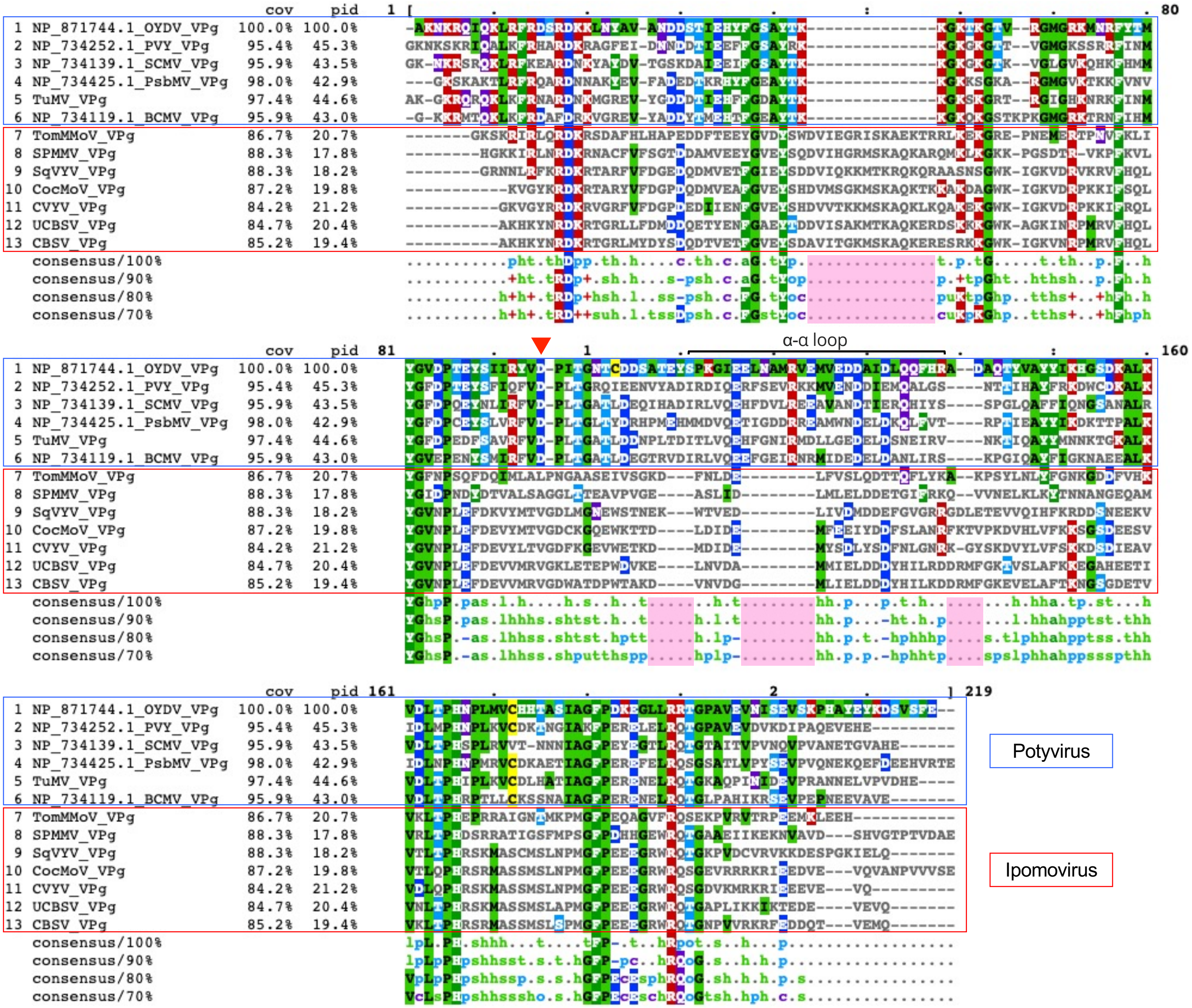


| #symbols => | color | #comment |
|---|---|---|
| . -> | dark-gray | #unconserved |
| + -> | bright-red | #positive charge |
| - -> | bright-blue | #negative charge |
| a -> | dark-green | #aromatic |
| c -> | purple | #charged |
| h -> | bright-green | #hydrophobic |
| l -> | bright-green | #aliphatic |
| o -> | dull-blue | #alcohol |
| p -> | dull-blue | #polar |
| s -> | bright-green | #small |
| t -> | bright-green | #turnlike |
| u -> | bright-green | #tiny |

Fig. S7 Alignment of *ipomovirus* and *potyvirus* VPg's show significant divergence in AA sequences

Representative potyvirus VPg sequences were aligned against all ipomovirus VPg sequences with the Clustal webtool and viewed through MView (Madeira *et al*., 2024). A conserved aspartate in potyvirus VPg proteins required for TuMV VPg interaction with *Arabidopsis* eIF(iso)4E is marked with a red triangle (Leonard *et al*., 2000). Pink highlighted regions of the consensus tracks indicate regions of sequence present only in one genus. The potyvirus α-α loop motif demonstrated to be the eIF4E interaction domain of PVY VPg is denoted by a horizontal bracket.

```
nCBP-2              MEITEKKDTENNTNNSNNNAQTTLDSASLENIDKEAEE-----------RQARDLKAGLH
nCBP-1              MEITE-EETENNNYSNNNARTASSP----DNIDKVAEE-----------RQARDLKAGLH
eIF4E               -MAAE-EPLKSTTEETPNPNLNSNPRAQDDVNDDEPEEGEIVGDEESSAKKSSAVTYQPH
eIF(iso)4E-1        -MATE-TATEGSATEATATGV------------EKPLQ---------------------H
eIF(iso)4E-2        -MASE-TAIEGTATEATASGA------------EQPQQ---------------------H
                       :*    :..  .                  . . :                     *

nCBP-2              PLKHKFVFWYTRRTPGVRTQTSYEDNIKKIVEFSTVEGFWVCYCHLARPSSLPSPTDLHL
nCBP-1              PLKHKFVFLYTRRTPGVRTQTSYEENIKKIMEFSTVEGFWVCYCHLARPSSLPIPTDLHL
eIF4E               PLEHQWTFWFDNPTAK-SKQATWGSSMRSIYTFATVEEFWSIYNNIHHPSKLAVGADFHC
eIF(iso)4E-1        KLERKWTFWFDNQSKP-KQGAAWGSSLRKVYTFDTVEEFWCLYDQIFKPSKLPGNADFHL
eIF(iso)4E-2        KLERKWTFWFDNQSKP-KQGAAWGTSLRKIYTFDTVEEFWCLYDQIFKPSKLPGNADFHL
                     *:.::.* : . :      :::  .:...: * *** **  * :: .**.*.  :*:*

nCBP-2              FKEGIRPLWEDSANSNGGKWIIRFKK-VVSGRFWEDMVLALVGDQLDYGDNICGAVLSIR
nCBP-1              FKEGIRPLWEDSANSNGGKWIIRFKK-AVSGRFWEDLVLALVGDQLGYSDNVCGAVLSIR
eIF4E               FKYKIEPKWEDPVCANGGKWTVTFGR-GKSDTSWLYTLLAMIGEQFDHGDEICGAVVNVR
eIF(iso)4E-1        FKAGVEPKWEDPECANGGKWSVTCGRKAILDTIWLETLMALIGEQFDEADEICGVVASVR
eIF(iso)4E-2        FKAGIEPKWEDPECAGGGKWSVTCGRKAILDTMWLETLMALIGEQFDEADEICGVVASVR
                    **  : * ***.  :.**** :   .    .  *   ::*::*:*:. .*::**.* .:*

nCBP-2              FNEDILSVWNRNSSDHQAVMALRDSIKRHLKLPHSYVMEYKPHDASLRDNSSYRNTWLRG
nCBP-1              FNEDILSIWNRNASDHQAVMALRDSLKRHLKLPHNYVMEYKPHDASLRDNSSYRNTWLRG
eIF4E               IKQEKIALWTKNASNEAAQLSIGKQWKEFLDYNDTIG--FIFHEDAKKLDRGAKNRYTI-
eIF(iso)4E-1        QRQDKLALWTKTATNEAAQMSIGKKWKEIIDVTEKIT--YSFHDDSRR-ERSVKGRYNV-
eIF(iso)4E-2        QRQDKLALWTKTATNEAAQMGIGKKWKEIIDVTDKIT--YSFHDDSRR-ERSAKGRYNV-
                     .:: :::*...:::  * :.: .. *  :.   .    :  *: : . : . .. :
```

| Residue | Color | Property |
|---|---|---|
| AVFPMILW | RED | Small (small+ hydrophobic (incl.aromatic -Y)) |
| DE | BLUE | Acidic |
| RK | MAGENTA | Basic - H |
| STYHCNGQ | GREEN | Hydroxyl + sulfhydryl + amine + G |
| Others | GRAY | Unusual amino/imino acids etc |

* fully conserved residue

: convservation between groups of strongly similar properties
STA NEQK NHQK NDEQ QHRK MILV MILF HY FYW

. conservation between groups of weakly similar properties
CSA ATV SAG STNK STPA SGND SNDEQK NDEQHK NEQHRK FVLIM HFY

Fig. S8 Alignment of cassava eIF4E-family proteins

Alignment was created and visualized with the Clustal Omega webserver (10.1002/pro.3290). nCBP-2 residues found to be important for CBSV VPg interaction and also conserved in at least one other family member are boxed in red.

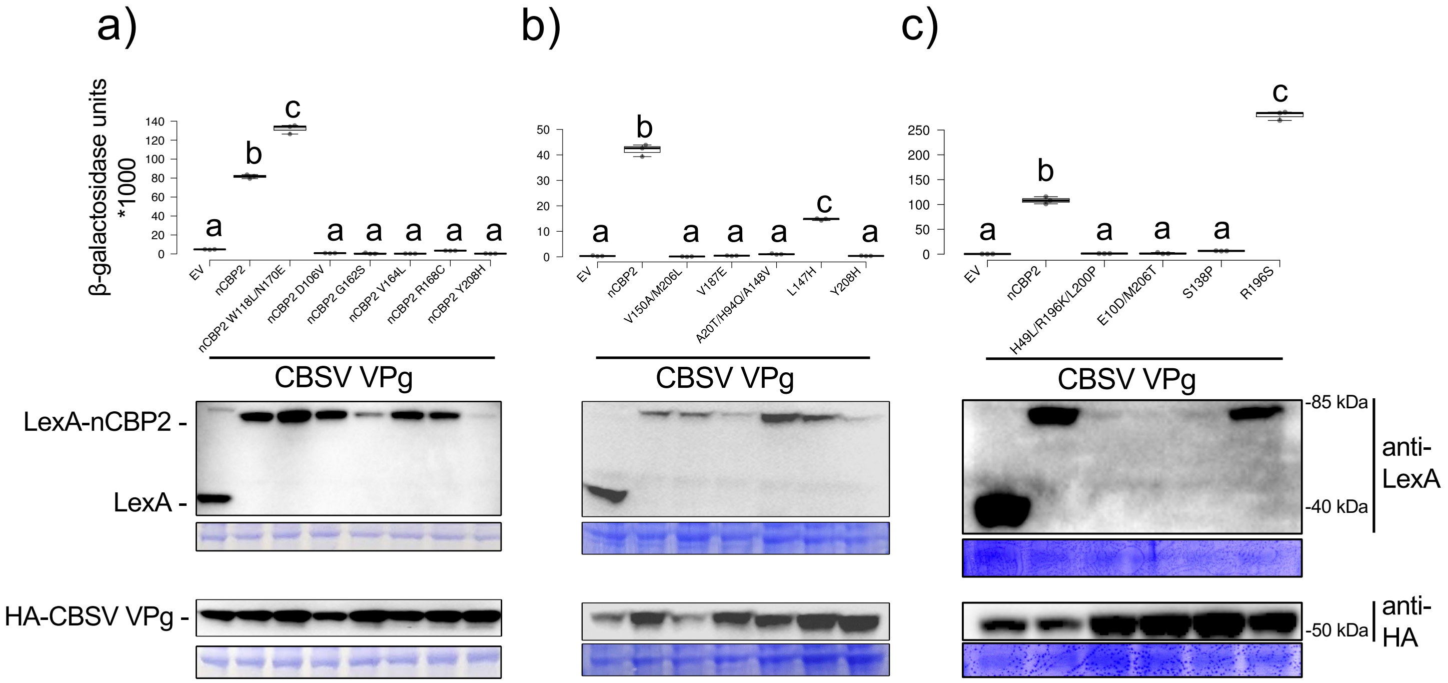


Fig. S9 Quantitative yeast two-hybrid assay confirms loss-of-affinity screen results

β-galactosidase activity was determined via a permeabilized cell assay for indirect measurement of of CBSV VPg interaction with cassava nCBP-2 variants. Assays were performed after inducing VPg expression by growing yeast for five hours in dropout media supplemented with galactose. Statistical differences were detected using one-way ANOVA and post hoc Tukey's HSD. LexA-nCBP-2 and HA-CBSV VPg expression in cells used for yeast two-hybrid was examined by western blot. Images of coomassie stained membranes are presented immediately below blot images.

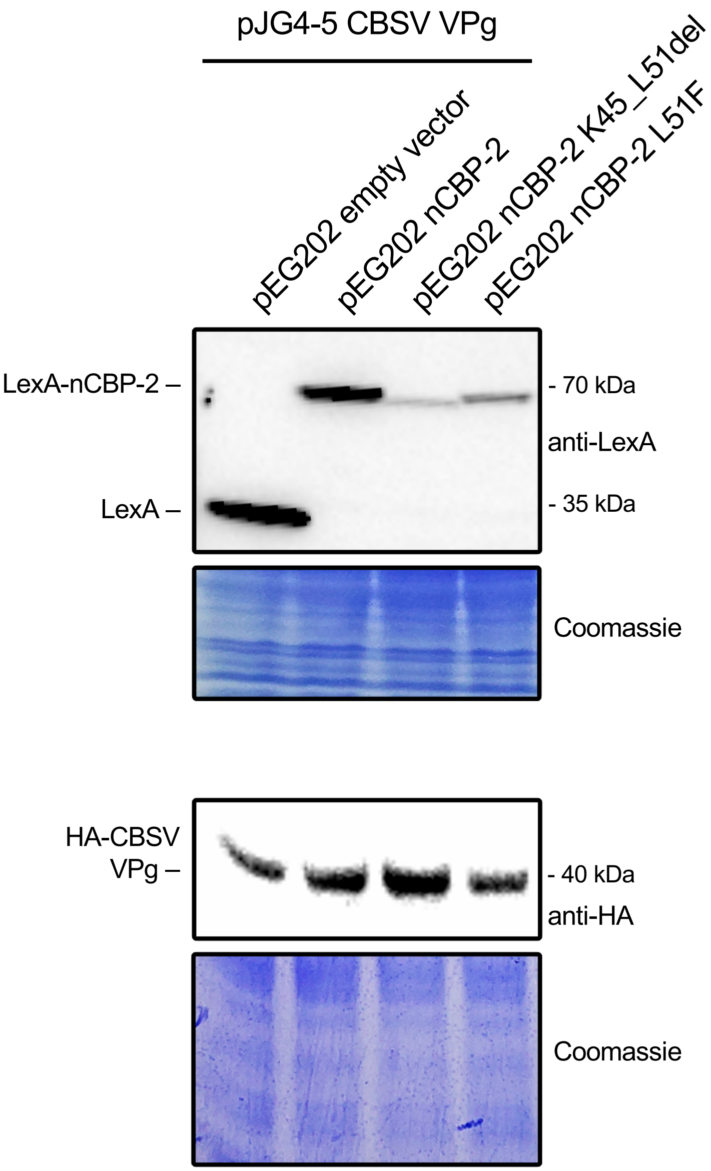


Fig. S10 Western blot analysis of yeast strains used for yeast two-hybrid experiments. A segment of coomassie-stained membrane is presented under corresponding western blots for assessing equal sample loading.

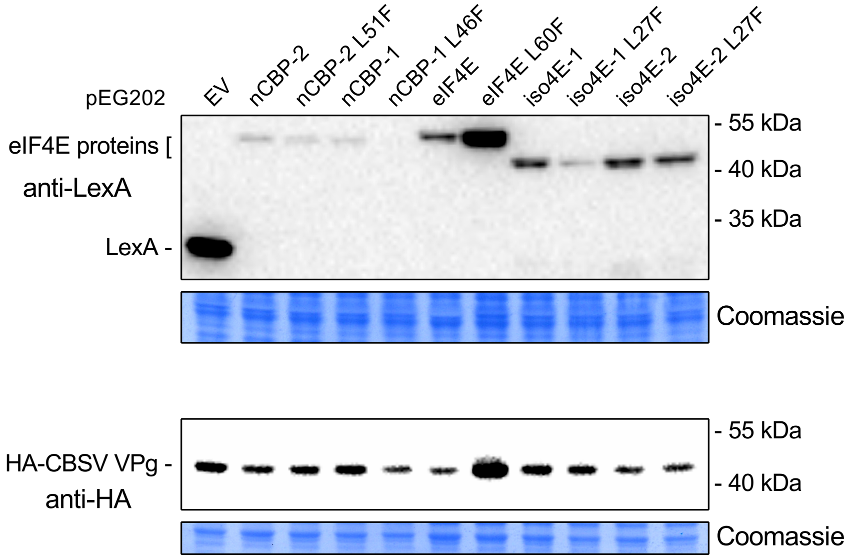


Fig. S11 Accumulation of eIF4E-family proteins and CBSV VPg in yeast used for quantitative yeast two-hybrid analysis

Western blot analysis of protein extracts from yeast used in the quantitative yeast two-hybrid experiment shown in figure 7. Coomassie stained membranes are shown under western panels for loading control.

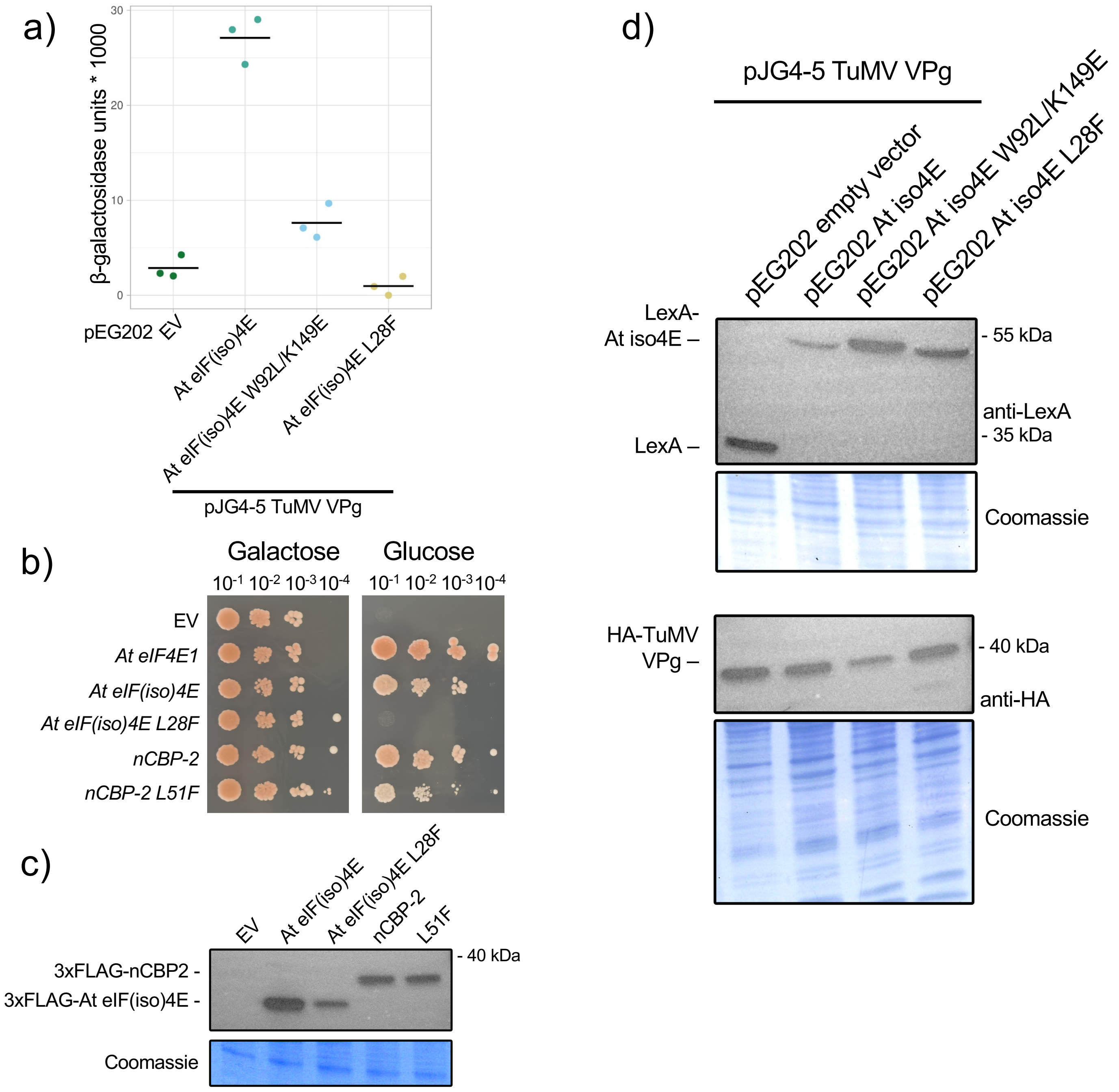


Fig. S12 Effects of mutating the HPL motif leucine in *Arabidopsis* eIF(iso)4E

a) Quantitative yeast two-hybrid analysis of TuMV VPg interaction with wild type *Arabidopsis* eIF(iso)4E alongside W92L/K149E and L28F mutants. Assays were performed after inducing VPg expression for five hours.
b) T93C *eif4e* yeast complementation assay with wild-type and mutant *eIF4E*-family genes from *Arabidopsis* and cassava.
c) Western blot analysis of eIF(iso)4E and nCBP-2 variants from cells used in (b).
d) Western blot analysis of protein accumulation in cells used in (a).
